# *Identifying inputs to visual projection neurons in* Drosophila *lobula by analyzing connectomic data*

**DOI:** 10.1101/2022.02.02.478876

**Authors:** Ryosuke Tanaka, Damon A. Clark

## Abstract

Electron microscopy-based connectomes provide important insights into how visual circuitry of fruit fly *Drosophila* computes various visual features, guiding and complementing behavioral and physiological studies. However, connectomic analyses of lobula, a putative center of object-like feature detection, remains underdeveloped, largely because of incomplete data on the inputs to the brain region. Here, we attempted to map the columnar inputs into the *Drosophila* lobula neuropil by performing connectivity- and morphology-based clustering on a densely reconstructed connectome dataset. While the dataset mostly lacked visual neuropils other than lobula, which would normally help identify inputs to lobula, our clustering analysis successfully extracted clusters of cells with homogeneous connectivity and morphology, likely representing genuine cell types. We were able to draw a correspondence between the resulting clusters and previously identified cell types, revealing previously undocumented connectivity between lobula input and output neurons. While future, more complete connectomic reconstructions are necessary to verify the results presented here, they can serve as a useful basis for formulating hypotheses on mechanisms of visual feature detection in lobula.

## Introduction

During the past decade, studies of the visual system of the fruit fly *Drosophila melanogaster* have generated an exquisitely detailed picture of how various visual computations are achieved by circuits of neurons (Cheong et al., 2020; de Andres-Bragado and Sprecher, 2019; Schnaitmann et al., 2018; Yang and Clandinin, 2018). In addition to the relative numerical simplicity of the fly brain (Raji and Potter, 2021) and its sophisticated genetic tools to monitor and manipulate specific cell types (Jenett et al., 2012; Kazama, 2015; Kitamoto, 2001; Pfeiffer et al., 2010; Simpson and Looger, 2018; Tirian and Dickson, 2017), a major contributor to this rapid progress has been electron microscopy (EM) based connectomes (Meinertzhagen, 2018, 2016). For example, dense EM reconstruction of circuitry surrounding the elementary motion detecting neurons in the *Drosophila* brain, T4 and T5 (Shinomiya et al., 2019, 2014; Takemura et al., 2015, 2017, 2013), have guided functional studies by providing strong constraints on neural computations in T4 and T5 (Agrochao et al., 2020; Behnia et al., 2014; Borst, 2018; Kohn et al., 2021; Ramos-Traslosheros and Silies, 2021; Strother et al., 2017; Zavatone-Veth et al., 2020), as well as by discovering previously undocumented circuit elements (Ammer et al., 2015; Behnia et al., 2014; Meier and Borst, 2019; Serbe et al., 2016; Strother et al., 2017). Similarly, EM reconstruction of early visual neuropils led to the discovery of a novel pathway for color vision (Takemura et al., 2008), which was later functionally confirmed to be contributing to the spectrally-sensitive behaviors (Gao et al., 2008).

In addition to motion and color, the visual system of flies is equipped with neurons tuned to a variety of visual features. Recent studies have shown that a series of columnar output neurons of lobula (**Fig. 1A**) collectively named lobula columner (LC) and lobula plate-lobula columnar (LPLC) neurons are tuned to visual features resembling conspecifics and predators, and their activity can induce a variety of behavioral responses, depending on types (Ache et al., 2019; Keleş and Frye, 2017; Klapoetke et al., 2017; Ribeiro et al., 2018; Städele et al., 2020; Sten et al., 2021; Tanaka and Clark, 2021, 2020; von Reyn et al., 2017; Wu et al., 2016). Several of these neurons are thought to achieve their selective visual tuning by pooling the feature selectivity of their presynaptic partners (Keleş et al., 2020; Klapoetke et al., 2017; Tanaka and Clark, 2021, 2020). However, the presynaptic partners of these LC and LPLC neurons remain largely unknown. This is due to the lack of connectomic reconstruction encompassing lobula and its upstream neuropils. For example, a recent connectomic dataset used to analyze the motion detection circuitry included most of the visual system, but the reconstruction was focused on T4 and T5 and their inputs, leaving out the most of lobula circuitry (Shinomiya et al., 2019). Similarly, community-driven tracing and proofreading effort of the full adult fly brain (FAFB) dataset (Zheng et al., 2018) is still in progress. While the densely-reconstructed hemibrain dataset contains almost the entire lobula, since it lacks almost entire medulla, neurons providing inputs into lobula (e. g., transmedullar (Tm) neurons) are fragmented and unlabeled (Scheffer et al., 2020).

**Figure 1.**
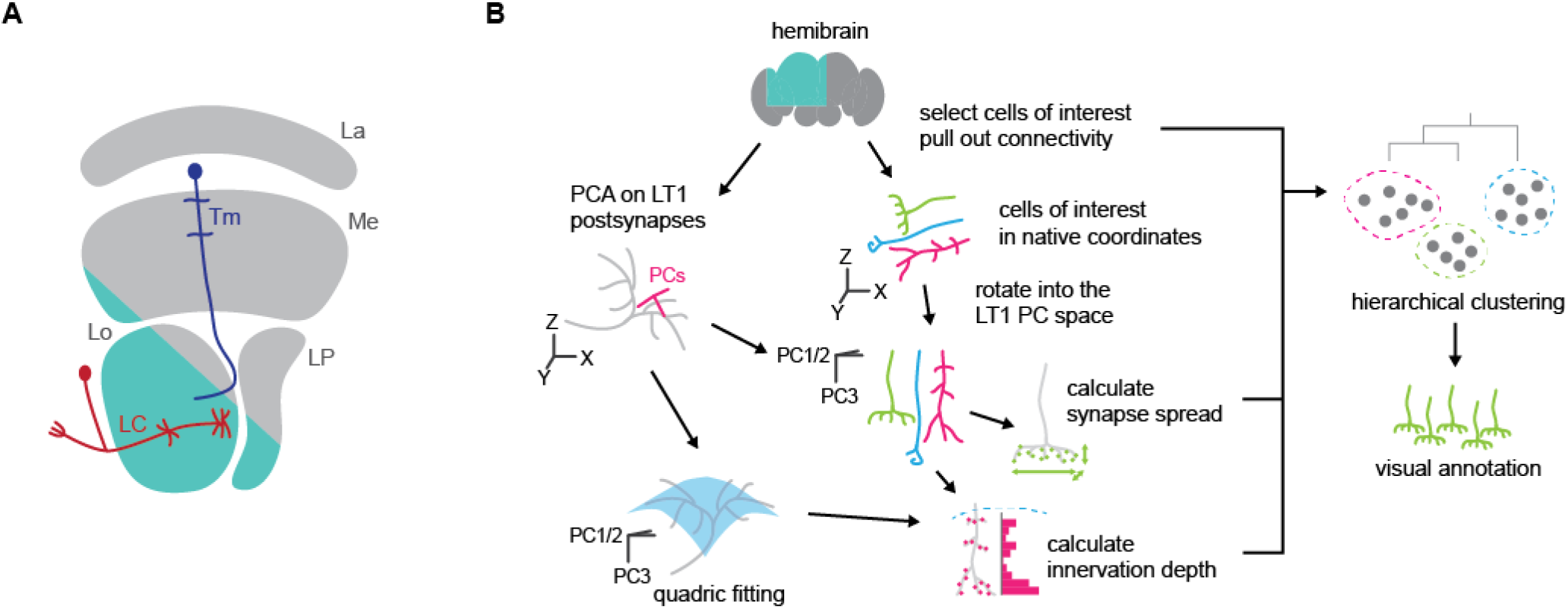
An overview of the present study. (A) A schematic of the *Drosophila* optic lobe. The optic lobe consists of four neuropils, namely, lamina (La), medulla (Me), lobula (Lo), and lobula plate (LP). Lobula houses projection neurons such as lobula columnar (LC) and lobula tangential (LT), which send axon terminals to the central brain and are thought to be responsible for detecting various visual features for approach and avoidance behaviors. Neurons connecting medulla and lobula, such as transmedullar (Tm) neurons, are likely providing major synaptic inputs into LCs and LTs, but their connectivity remain largely unknown. The teal shaded region roughly indicates the volume reconstructed in the hemibrain dataset. (B) A schematic of the analysis pipeline. Small lobula intrinsic neurons in the hemibrain dataset were selected as cells of interest, and their connectivity to labeled downstream neurons were tabulated. The candidate cells were then rotated into the principal component (PC) space of LT1 postsynapses, which we used as a proxy for layer 2 of lobula, from which we derived normal and tangential axes of the lobula layers. We then quantified the morphology of the cells of interest by synapse spread in the PC space and by the histogram of innervation depth relative to a parabolic surface fit to the LT1 synapses. Finally, we applied hierarchical clustering to the concatenated feature matrix, and visually annotated the resulting clusters.

However, recent studies have shown that close inspection of these fragmented axon terminals of input neurons into lobula in the hemibrain dataset can still offer potentially useful information about the lobula circuitry (Tanaka and Clark, 2021, 2020). In the present study, we perform clustering analyses on those fragmented terminals based on their connectivity and morphology, defining putative inputs cell types for intrinsic and projection neurons of lobula, including LC and LPLC neurons. This analysis comes with the caveat that it is difficult to conclusively identify cell types based on their axonal morphology alone and thus needs to be complemented by a more comprehensive connectome encompassing the entire optic lobe. Even so, the results presented here provide a previously unknown putative connectivity to physiologists and psychophysicists interested in visual processing in lobula.

## Methods

Code to run the analyses performed in this study is available on a GitHub repository (https://github.com/ClarkLabCode/LobulaClustering). The goal of present analysis is to categorize unlabeled putative columnar neurons in lobula into cell types, and examine their connectivity to downstream neurons. Typically, a columnar visual neuron type consists of many cells sharing stereotypic connectivity and morphology tiling the visual space. Thus, the problem boils down to grouping neurons by their individual connectivity and morphology while discounting their exact spatial location. To this end, we performed an agglomerative clustering on their connectivity and morphology. A schematic of the analysis pipeline is shown in **Figure 1B**.

### Inclusion criteria

Cells without cell type assigned in the hemibrain 1.2.1 dataset that had synapses only in the lobula and had more than 50 and less than 500 total synapses were selected. We refer to these cells as “cells of interest”. 7,465 cells of interest met the above criteria and were incorporated to the analyses.

### Connectivity

For each cell of interest, we counted the number of synapses it has onto labeled cell types in lobula. Connections to multiple cells of a single cell type were summed together to generate a single feature, rather than being treated separately. This was intended to facilitate grouping of cells belonging to the same cell types with a shared pattern of connectivity regardless of their retinotopic location. A total 237 labeled cell types were found to be downstream of at least one of the cells of interest in the present analysis.

### Morphology

To summarize features of the axonal morphology of the cells of interest, we calculated (1) the standard deviation of synapse position along the two tangential axes and the normal axis of the lobula layers, as well as (2) the histogram of synapses along the depth of lobula. The native coordinate system of the hemibrain dataset is a Cartesian system where X points left, Y points anteriorly, and Z points ventrally (Scheffer et al., 2020). To identify the normal and tangential axes of the lobula layers, we first performed principal component analysis (PCA) on the postsynapses of LT1 neurons. LT1 is a tangential neuron with wide, dense, monostratified dendrites in the layer 2 of lobula (Fischbach and Dittrich, 1989). The hemibrain dataset contains three LT1 neurons, each covering the entire tangential extent of the lobula. The resulted first two principal component (PC) axes were approximately aligned to the ventrolateral and anteroposteior axes of lobula, respectively, and the third PC axis was normal to that plane. Then, for each cell of interest, we calculated the standard deviations of the distributions of its presynapses along the three PC axes, which quantified tangential extents and vertical diffuseness of its axon terminal. We refer to these three features as “synapse spread” hereafter.

Next, we aimed to calculate the depth of each synapse along the layers of lobula. Since the layers of the lobula are curved, the raw position of the synapses along the third PC axis do not accurately correspond to their layer affiliation. To solve this issue, we first fit a parabolic surface

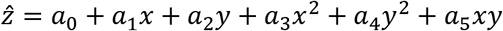

to the postsynapses of the LT1 neurons with the least-square method, where *x* and *y* represent position along the first and second PC axes (μm), *ẑ* represents the predicted position along the third PC axis (μm), and *a_i_* are coefficients. The fit coefficients were as follows: a_0_ = −5.21, a_1_ = 4.04 × 10^−3^, a_2_ = 4.04 × 10^−3^, a_3_ = 3.17 × 10^−3^, a_4_ = 7.65 × 10^−4^, a_5_ = −1.22 × 10^−3^. The goodness of fit was R^2^ = 0.744. We then calculated the depth of each synapse relative to the layer 2 as *z* – *ẑ*, where *z* represents the true position of the synapse along the third PC axis. Note that this approximates the lobula layers to be parallel to each other, rather than concentric curved surfaces. Deviations from this approximation will be most severe far from the center of this PC coordinate system. For each cell of interest, we counted the number of presynapses in 5 μm wide bins ranging from −20 to +50 μm of relative depth, resulting in 14 features. This range covered the entirety of the lobula. We refer to these features as “innervation depth” in the following. Similar analyses of innervation depth have been performed previously on medulla neurons (Bausenwein et al., 1992; Zhao and Plaza, 2014). We validated that the synapse spread and innervation depth calculated this way appropriately reflect the neuronal morphology by applying the same analysis to postsynapses on the dendrites of multiple LC types with well characterized morphology (Wu et al., 2016).

### Agglomerative clustering on cells of interest

We concatenated the connectivity, synapse spread, and smoothed innervation depth feature matrices into a single feature matrix, and performed an agglomerative clustering with Ward’s variance minimization method (Ward, 1963) with SciPy (Virtanen et al., 2020). Before being concatenated, connectivity, synapse spread, and innervation depth matrices were respectively normalized by its total dispersion (sum of variances) and then weighted by the factors of 5, 1, and 3, respectively. This weighting reflected our relative confidence on the accuracy of the three feature sets. We set the number of clusters to be 40, which is approximately comparable to the number of transmedullar (Tm) and transumedullar Y (TmY) cell types reported in Fischbach & Dittrich (1989). The complete results of the clustering analysis are presented in **Supplementary File 1**.

### Annotation and visualization

Next, we visualized the cells of interest in each resultant clusters on the neuPrint website (Clements et al., 2020a) and compared their terminal morphology with existing anatomical literature (Fischbach and Dittrich, 1989) to determine cell type identity. We also performed a nonlinear dimensionality reduction on the concatenated feature matrix with UMAP (McInnes et al., 2018) for visualization.

### Quantification of connectivity between cells of interest and visual projection neurons

Lastly, we analyzed the connectivity between the 40 clusters of cells of interest and 21 LC and LPLC (Wu et al., 2016). For each LC or LPLC type, we calculated relative number of synapses they receive as a population from each of the 40 clusters. Additionally, an agglomerative clustering was performed on the relative connectivity matrix between the 40 clusters and 21 LC/LPLCs.

## Results

### The synapse spread and innervation depth features capture the dendritic morphology of LC neurons

To gain confidence in our quantification of cellular morphology, we first calculated innervation depth and synapse spread features for lobula postsynapses of 18 lobula columnar (LC) and lobula plate-lobula columnar (LPLC) projection neuron types with well characterized morphology (LC4, 6, 9, 11-13, 15-18, 20-22, 24-26, LPLC1 and 2) (Wu et al., 2016) (**Fig. 2**). **Figure 2A and B** show previously reported light microscopy-based innervation patterns of LCs and LPLCs (Wu et al., 2016) (**Fig. 2A**) side by side with their connectome-based innervation depth feature (**Fig. 2B**). The patterns of innervation captured by two methodologies agreed well. For example, the bistratified dendrites of LC4 in the layers 2 and 4 are clearly visible from the connectome-based innervation depth features (**Fig. 2A, B,** the leftmost column). The synapse spread of LCs and LPLCs along the two tangential and the normal axis of lobula are shown in **Figure 2C and D**. These features also captured well the known morphology of these neurons. For example, LC25 and 12, which respectively had large and small synapse spread along the both tangential dimensions (**Fig. 2C**) have been previously shown to have large and small isotropic dendrites (Wu et al., 2016 – Figure 6). LC22, which had large synapse spread along the long tangential axis of lobula but small spread along the short axis, indeed has anisotropic, narrow dendrites (Wu et al., 2016 – Figure 6). In a similar vein, along the normal axis, vertically diffuse neurons such as LPLC1 and LC15 (Wu et al., 2016 – Figure 5) had large synapse spread, whereas monostratified LC25 had small spread (**Fig. 2D**). Overall, these results indicate that the two morphological features we calculated accurately capture the morphology of neurites in the lobula neuropil.

**Figure 2.**
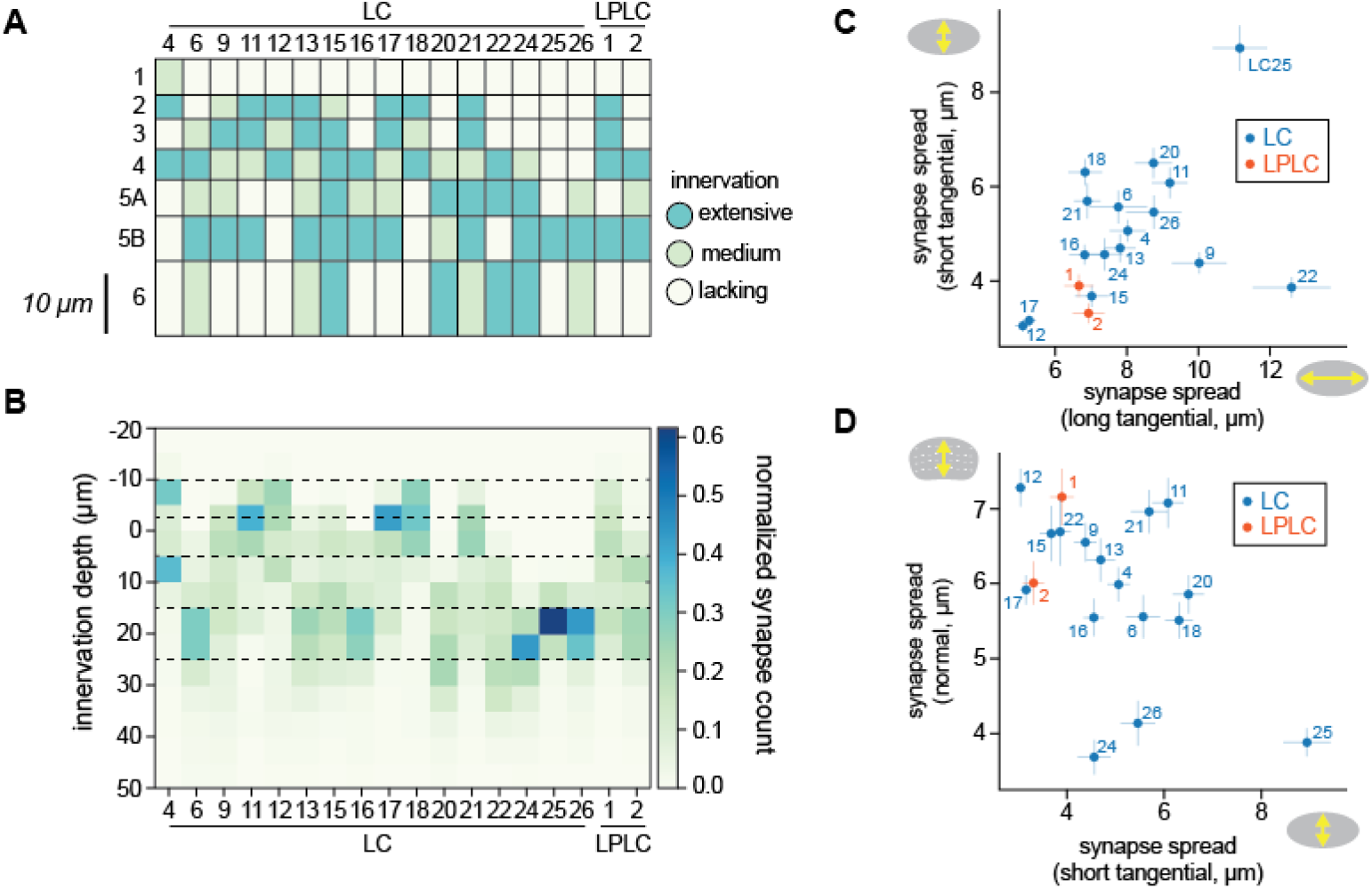
Validation of morphological features. (A) A diagram showing innervation patterns of LC and LPLC neurons in lobula, based on a previously published data (Wu et al., 2016 – Figure 6). (B) Innervation depth features of the identical set of LC and LPLC neurons as in (A). Innervation depth were averaged across all the cells in each cell type and then normalized within each type. The horizontal dotted lines indicate the boundaries between layers inferred by comparing (A) and (B). (C, D) Mean synapse spread of LCs and LPLCs along the long and short tangential axes and the normal axis, with standard error of the mean.

### Agglomerative clustering identifies clusters with homogeneous connectivity and morphology

The results of the clustering are shown in **Figures 3, 4** and **Table 1**. The clustering successfully extracted groups of cells with similar connectivity and morphology, as can be seen from the block-like appearance of the feature matrices (**Fig. 3B-D**). Additionally, we visualized the cells of interest on the two-dimensional nonlinear embedding by uniform manifold approximation (UMAP) (**Fig. 3E**). Overall, the embedding agreed well with the hierarchical clustering, since neighboring clusters are often neighbors in the UMAP space. This was not without exceptions, such as clusters 15, 16, 29, and 31, which had clear substructures on the UMAP embedding. **Table 1** summarizes the morphology, connectivity, and putative cell type affiliation of all identified clusters. Example clusters that we identified with previously documented types of neurons with reasonable confidence are shown in **Figure 4**. The input columnar cell types we identified most confidently included T3, T2, TmY9/Tm28, Tm20, and TmY11 (**Fig. 4A**). The **Appendix** contains detailed analyses of each cluster.

**Figure 3.**
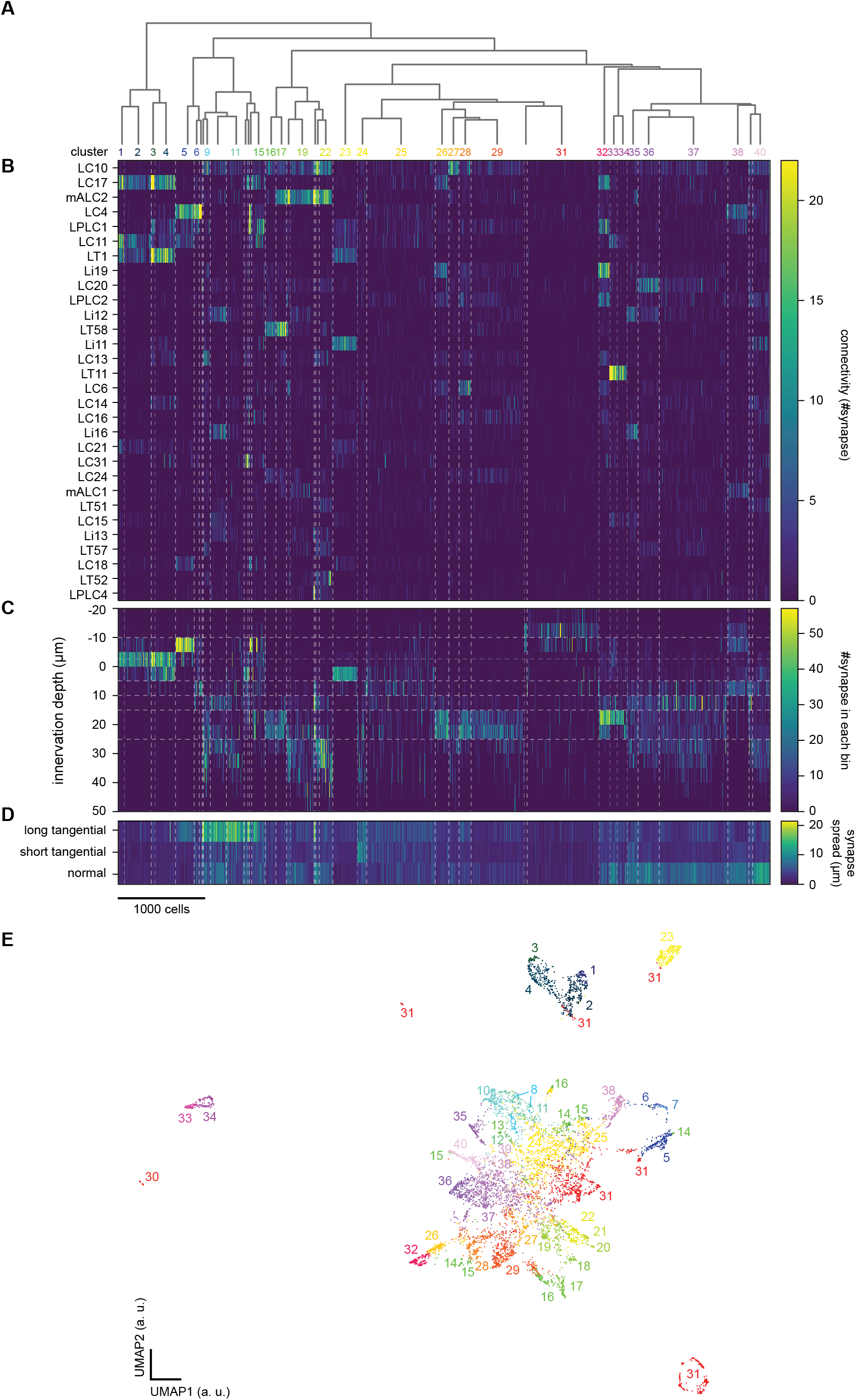
Agglomerative clustering of unlabeled lobula neurons. (A) The dendrogram of the clusters, with cluster labels. Only clusters with more than 50 cells are labeled for visibility. (B) Connectivity between the cells of interest, sorted by cluster affiliation, and the top 30 labeled lobula neuron types with the largest number of synapses from the cells of interest. The white dotted lines delineate clusters, throughout B-D. (C) Innervation depth of the cells of interest, sorted by cluster affiliation. The horizontal dotted lines indicate approximate layer boundaries, as in **Figure 2B**. (D) Synapse spread of the cells of interest along the tangential and normal axes of lobula. (E) The cells of interest on the UMAP embedding space, based on the normalized, weighted, and concatenated feature. Clustering was performed directly on the feature matrix, so that the UMAP embedding is simply for visualization. a. u.: arbitrary unit.

**Figure 4.**
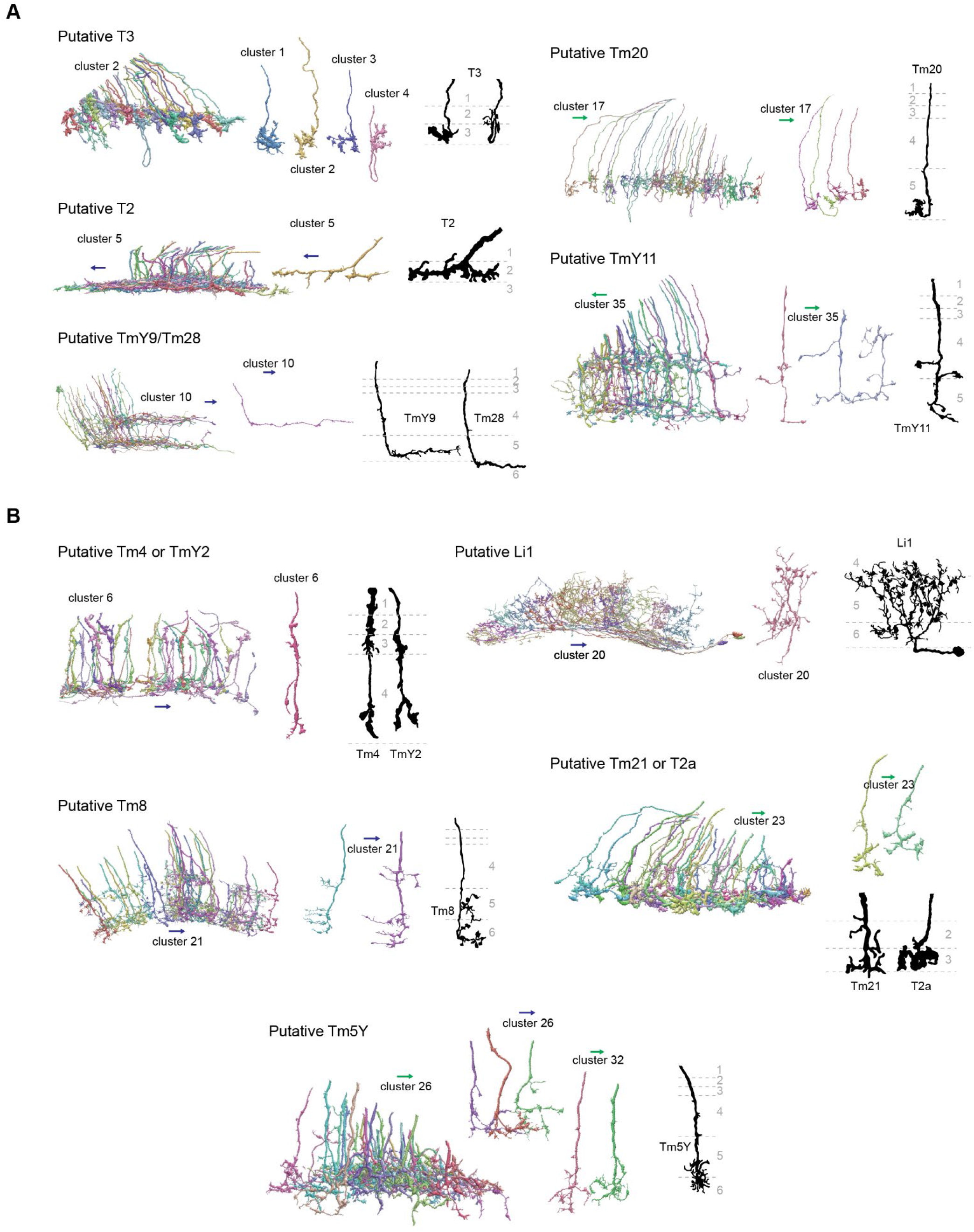
Example clusters with putative morphological matches. (A) Clusters morphologically identified with known cell types with the highest confidence, labeled as “A” in **Table 1**. For each putative cell type, (*left*) a population of reconstructed neurons in the corresponding clusters viewed tangentially, (*middle*) individual examples of cells in the clusters, and (*right*) Golgi staining from Fischbach & Dittrich (1989) are shown. (B) The same as (A), but for clusters with less confidence (labeled as “B” in **Table 1**). The green and blue arrows respectively point towards anterior and ventral directions.

**Figure 5.**
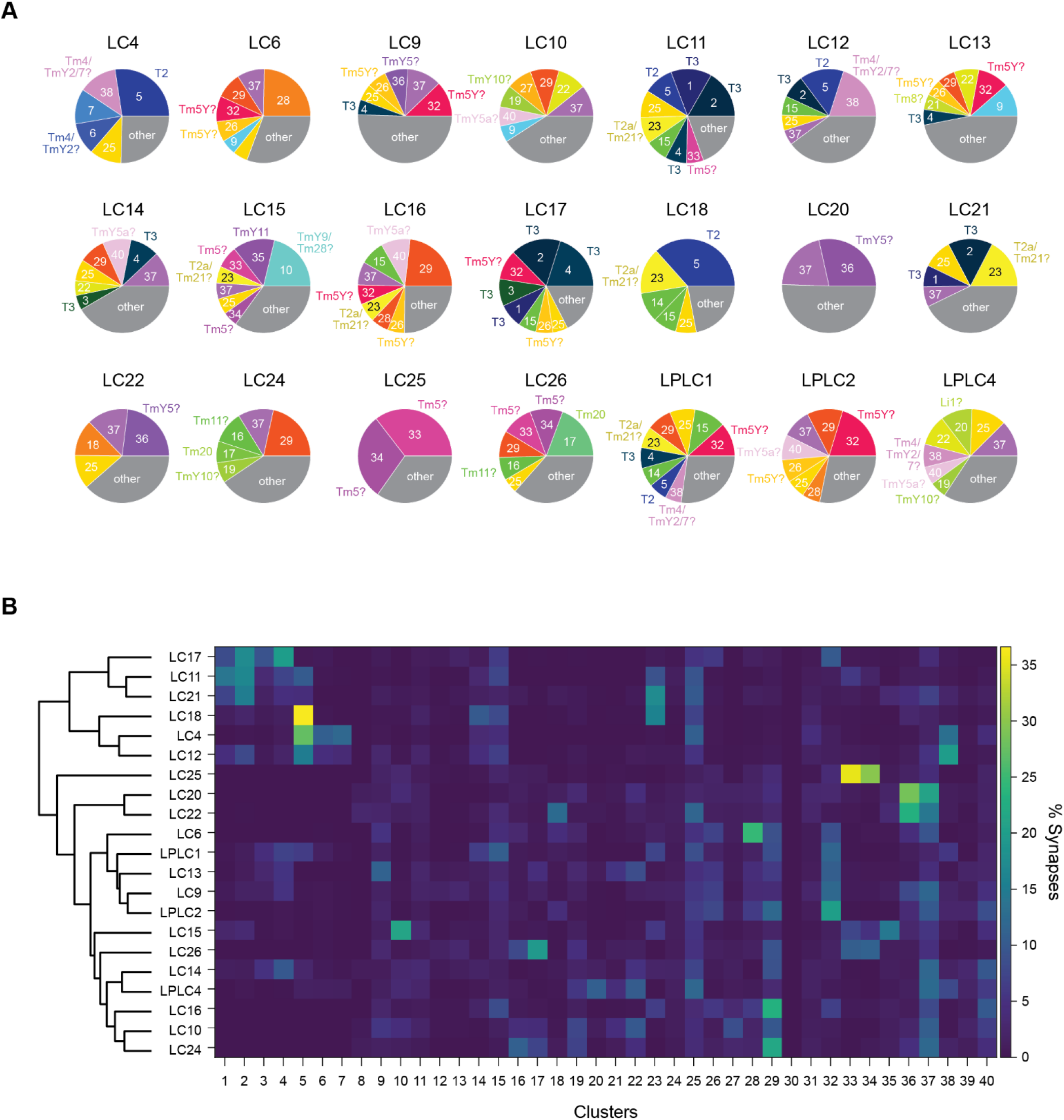
Inputs to columnar visual projection neurons. (A) Relative connectivity from the 40 clusters of the cells of interest and LC/LPLC neurons. Clusters with less than 5% of synaptic input from cells of interest were lumped into “others”. Putative cellular identities of clusters are also shown. Labels with question marks indicate confidence less than “B” in **Table 1**. (B) Relative connectivity from the 40 clusters to the 21 LC/LPLC types, with a dendrogram from the agglomerative clustering analysis of the LP/LPLC types. Connectivity is normalized for each LC/LPLC type, such that each row of the connectivity matrix adds up to 1.

**Table 1.** Summary of clustering on the cells of interest. The homogeneity of each cluster was ranked in three tiers: A – likely single cell type, B – several cell types, and C – mixture of many cell types. Cell types are labeled where identified, and a question mark is used in cases of cell types that appear homogeneous but previously undocumented. The confidence of cell-type identification was also ranked in three tiers (A, B, and C for highest to lowest confidence) or marked NA for clusters where we were unable to find cell types. See **Appendix** for the detailed analysis of each cluster.

| # | Homogeneity | Putative cell type | Confidence | Layers | Morphology | Connectivity | Related to |
| --- | --- | --- | --- | --- | --- | --- | --- |
| 1 | A | T3 | A | 2, 3 | small, crumpled, sometimes with loops | LC17, LC11, LT1 | #2-4 |
| 2 | A | T3 | A | 2, 3 | small, crumpled, sometimes with loops | LC17, LC11, LT1 | #1,3,4 |
| 3 | A | T3 | A | 2, 3 | small, crumpled, sometimes with loops | LC17, LC11, LT1, LPLC1 | #1,2,4 |
| 4 | A | T3 | A | 2, 3 | small, crumpled, sometimes with loops | LC17, LC11, LT1, LPLC1 | #1-3 |
| 5 | A | T2 | A | 2 | T shaped, spiny | LC4, LC11, LC18, Li15, LPLC1 |  |
| 6 | B | Tm4 or TmY2 | B | 2, 4 | small, vertical, Y-shaped ends, swelling in Lo2 | LC4, LPLC1 |  |
| 7 | C | mixture | NA | 2, 4 |  | LC4, LC12 |  |
| 8 | A | ? | NA | 6 | wide field, lobula intrinsic | LC10, LC20, mALC2, LT52, Li12 |  |
| 9 | A | ? | NA | 5B | anisotropic, bent dorsally | LC10, LC13, Li19, LC6, LT57 |  |
| 10 | B | TmY9 or Tm28 | A | 4, 5B | Linear, L-shaped, optional branching and layer-switching | LC10, Li16, Li12, LC15, Li13 |  |
| 11 | C | mixture | NA | 4-6 |  | LC10 |  |
| 12 | A | Tm19 | C | 3-5B | meandering, large secondary branch | LT87, LC31, LC17 |  |
| 13 | A | Tm19 | C | 3-5B | meandering, large secondary branch | LT87, LC31, LC17 |  |
| 14 | C | mixture | NA | 2-5A |  | LPLC1 | #5 |
| 15 | C | mixture | NA | 2-5B |  | LPLC1 |  |
| 16 | A | Tm11 | C | 5B | thin, knobby, posteriorly oriented | LT58, LC10, LC24 |  |
| 17 | A | Tm20 | A | 5B | thick, compact | LT58, mALC2, LC10, LC24 | #29 |
| 18 | A | ? | NA | 5B, 6 | short branches, knobby endings | mALC2, LPLC2, LT58, LC22, LC6 |  |
| 19 | A | TmY10 | C | 5A-6 | somewhat vertically diffuse, small protrusions in shallower layers | mALC2, LC10, mALC1, Li12 |  |
| 20 | A | Li1 | B | 4, 5A, 6 | Lobula intrinsic, bistratified, knobby | mALC2, LPLC4, LC10, Li13, LT36 |  |
| 21 | A | Tm8 | B | 5A, 6 | Bistratified, dorsally oriented multiple branches | mALC2, LC10, LT57, LC13, LT52 |  |
| 22 | B | ? | NA | 5A, 6 | thick, or thin and knobby (two cell types) | mALC2, LC10, LT52, LT63, LT51 |  |
| 23 | A | T2a or Tm21 | B | 3 | Monostratified, small, crumpled | Li11, LT1, LC11, LPLC1, LC21 |  |
| 24 | C | mixture | NA | diffuse |  | mALC2 |  |
| 25 | C | mixture | NA | diffuse |  | LC4 |  |
| 26 | A | Tm5Y | B | 5A/B | Thick, ventrally oriented, monostratified, small protrusions in shallower layers | Li19, LC17, LT79, LPLC2, LC6 | #32 |
| 27 | B | ? | NA | 5B | Thin, monostratified, knobby endings | LC10 | #29 |
| 28 | A | ? | NA | 5A/B | Thin, monostratified, knobby endings | LC6, LPLC2, LC10, Li19, LC16 |  |
| 29 | B | ? | NA | 5A/B |  | LC10, LC16 | #17, 27 |
| 30 | A | Tm9 | A | 1 | Small, crumpled | CT1 |  |
| 31 | C | mixture | NA | 1 | T5 and its inputs |  |  |
| 32 | A | Tm5Y | B | 5A/B | Thick, ventrally oriented, monostratified, small protrusions in shallower layers | Li19, LC17, LT79, LPLC2, LPLC1 | #26 |
| 33 | A | Tm5 | B | 5A/B | Straight, small protrusion in a shallower layer | LT11, LC25, LC11, LC15 | #34 |
| 34 | A | Tm5 | B | 5A/B | Straight, small protrusion in a shallower layer | LT11, LC25, LC11, LC15 | #33 |
| 35 | A | TmY11 | A | 4-6 | Perpendicular branches along the short axis | Li16, Li12, LC15, LC10, Li19 |  |
| 36 | A | TmY5 | C | 4-6 | Many short protrusions | LC20, Li12, LC22, LC10, Li19 |  |
| 37 | C | mixture | NA | 4-6 |  | LC10, LC20 | #32-36 |
| 38 | B | Tm4 or TmY2 and TmY7 | B | 1-5A | Short, vertical, Y-shaped ending + knobby | LC4, Li17, mALC1, LPLC1, LC12 | #6 |
| 39 | A | ? | NA | 4-6 | Branching, knobby endings | Li14, LC10, PS179, LPLC2, LC14 |  |
| 40 | A | TmY5a | C | 3-6 | Vertically diffuse, bistratified, minor protrusions<br>in between | LC10, Li11, LPLC2, LT51, LC4 |  |

### Inputs into columnar visual projection neurons

To gain insight into the visual computations of columnar visual projection neurons in lobula, we calculated connectivity between the 40 clusters of the cells of interest and 21 LC and LPLC neurons (Wu et al., 2016). **Figure 5A** summarizes major presynaptic clusters for each of the 21 LC/LPLC type, with putative cellular identities of the clusters labeled. **Figure 5B** also shows relative connectivity between 40 clusters and the 21 LC/LPLC neurons, but with a dendrogram from the agglomerative clustering analysis. The most fundamental divide in the dendrogram was between “shallow” LCs with prominent Lo2/3 innervation (LC4, 11, 12,17, 18, 21) (**Fig. 2A, B**) and the rest. Among the 6 shallow LC types, LC11, 17, and 21 received more inputs from T3 (clusters 1-4), whereas LC4, 12, and 18 received more inputs from T2 (cluster 5). Among the remaining cell types, LC25 stood out for its almost exclusive connectivity to Tm5 neurons. LC20 and LC22 also had a relatively distinct connectivity pattern, with prominent connection to cluster 36 (putative TmY5). The remaining cell types were divided into two branches, one of which included LC6, 9, 13 and LPLC1, 2, and the other including LC10, 14, 15, 16, 24, 26, and LPLC4. The former branch stood out for their shared connectivity to clusters 26 and 32, putative Tm5Y. The clusters shared by the latter branch included clusters 19 (putative TmY10) and 40 (putative TmY5a). Additionally, we found some clusters that provide substantial inputs to only single LC/LPLC type analyzed here. For example, LC15 was the only cell type that received more than 5% of its inputs from cluster 10 (putative TmY9/Tm28). Similarly, LC6 was the unique target of cluster 28 and LC22 was the unique target of cluster 18. We could not find morphological matches for neither cluster 18 or 28, but they were morphologically homogeneous, likely representing genuine cell types.

## Discussion

Lobula is one of the deepest neuropils in the fly visual system and is thought to detect behaviorally relevant visual features and send outputs to the central brain (Mu et al., 2012; Otsuna and Ito, 2006; Wu et al., 2016). While a number of cell types providing inputs to and sending outputs from lobula have been characterized, connectivity among them — critical information to understanding the computations taking place in lobula — have been largely missing. In the present study, we attempted to fill this gap by performing connectivity- and morphology-based clustering on fragmented lobula axon terminals in the hemibrain dataset (Scheffer et al., 2020). Among the 40 clusters, we were able to find some morphologically matching cell types for 24 clusters, with varying degrees of confidence (**Table 1**). The cell types we identified with most confidence were: T3 (clusters 1 through 4), T2 (cluster 5), TmY29 and/or Tm28 (cluster 10), Tm20 (cluster 17), Tm9 (cluster 30), and TmY11 (cluster 35). These were mostly cell types that had distinctive terminal morphology in lobula (Fischbach and Dittrich, 1989). The other identifications — Tm4 and/or TmY2 (clusters 6 and 38), Tm19 (clusters 12, 13), Tm11 (cluster 16), TmY10 (cluster 19), Li1 (cluster 20), Tm8 (cluster 21), T2a and/or Tm21 (cluster 23), Tm5Y (clusters 26 and 32), Tm5Y (cluster 32), Tm5 (clusters 33 and 34), TmY5 (cluster 36), TmY5a (cluster 40) — were more speculative.

The current dataset lacks the information required to definitively identify cell types: the innervation patterns in medulla and lobula plate. Future connectomic reconstructions encompassing the entire optic lobe will be necessary to unambiguously identify the putative cell types identified here. This can be in principle done by tracing in the full adult fly brain (FAFB) electron microscopy volume (Zheng et al., 2018) on the FlyWire website (Dorkenwald et al., 2021). An alternative approach is to use molecular tools to map connectivity among genetically defined cell types, such as trans-synaptic GFP reconstitution (GRASP) (Feinberg et al., 2008; Shearin et al., 2018), trans-Tango (Talay et al., 2017), and optogenetics (Simpson and Looger, 2018). This approach requires generating or identifying selective transgenic drivers to target new Tm and TmY neurons, and will be required to define functional connectivity, rather than anatomical connectivity. Neuroanatomical tools such as color depth maximum intensity projection mask search (Clements et al., 2020b; Otsuna et al., 2018) as well as marker gene identification with single cell RNA sequencing approaches (Özel et al., 2020) will be useful to this end. We found clusters consisting of apparently homogeneous cells whose morphology did not match any previously reported Tm or TmY cells (Fischbach and Dittrich, 1989) (clusters 8, 9, 18, 22, 27-29, 39), likely representing new cell types.

While most of the putative cell types identified here have never been studied physiologically, there are some exceptions. For example, Tm4 (clusters 6/38) and Tm9 (cluster 30) are OFF cells presynaptic to T5 (Serbe et al., 2016; Shinomiya et al., 2014). Tm5 (clusters 33/34) and Tm20 (cluster 17) are spectrally sensitive cells downstream of the inner photoreceptors (Gao et al., 2008; Lin et al., 2016; Melnattur et al., 2014). T2 (cluster 5) and T3 (clusters 1-4) are known LC/LPLC inputs with selectivity for small moving objects (Keleş et al., 2020; Tanaka and Clark, 2021, 2020). Interestingly, the spectrally sensitive Tm neurons and object selective neurons appeared to converge on several LC neurons. For example, both LC11 and LC15 received sizable inputs from putative T3 and Tm5, suggesting that these cell types may prefer objects of specific colors. For lobula input neurons without known physiology, their connectivity to previously characterized LC/LPLCs can hint at their function. For example, cluster 32 neurons (putative Tm5Y) provide sizable inputs to LC6, 9, 13 and LPLC1, 2. LC6 (Morimoto et al., 2020; Wu et al., 2016), LPLC1 (Tanaka and Clark, 2021), and LPLC2 (Ache et al., 2019; Klapoetke et al., 2017) are sensitive to looming stimuli, suggesting that Tm5Y might play an important role in loom detection. Another example is cluster 10 neurons (TmY9/Tm28), which selectively connect to LC15. LC15 respond preferentially to moving, narrow vertical bars (Städele et al., 2020). The highly anisotropic dendrites of TmY9/Tm28 oriented along the dorsoventral axis (Fischbach and Dittrich, 1989) could be well suited to pool excitation along the vertical visual axis to detect long bars. Overall, the analysis presented here advances our knowledge of lobula connectivity and generates testable hypotheses for how lobula visual projection neurons become selective for specific visual features.

## Acknowledgements

RT was supported by the Takenaka Foundation and the Gruber Foundation. DAC and this project were supported by NIH R01EY026555, NIH R01NS121773, NIH P30EY026878. RT and DAC conceived the study. RT performed the analysis and annotation. RT and DAC wrote the paper. Authors declare no competing interests.

## Appendix Detailed characterization of the 40 clusters

The following describes the detailed morphological analyses of the 40 clusters of the cells of interest resulting from the agglomerative clustering (**Fig. 3**), on which putative cell type identification in **Table 1** and **Figs. 4, 5** are based.

### Clusters 1 through 4

Clusters 1 through 4 were highly distinct from the other clusters, as can be seen from the dendrogram (**Fig. 3A**) and the UMAP embedding (**Fig. 3E**). Cells in these clusters had small monostratified axon terminals slightly elongated along the long tangential axis (synapse spread of about 2.5 micron along the long axis, and 1.5 micron along the other axes), which innervated the boundary between layers 2 and 3 of lobula (i. e., Lo2 and Lo3) (**Fig. A1A-C**). The major synaptic targets of these clusters were LC17 (13.7, 6.4, 25.0, and 9.9 synapses/cell for clusters 1, 2, 3, and 4, respectively), LC11 (15.8, 4.2, 6.3, and 2.5 synapses/cell for clusters 1, 2, 3, and 4, respectively), LT1 (8.1, 3.1, 22.4, and 12.3 synapses/cell for clusters 1, 2, 3, and 4, respectively). LPLC1 was also a major target for clusters 3 and 4 (8.7 and 2.5 synapses/cell for clusters 3 and 4, respectively). Visually, the four clusters appeared homogeneous, consisting of cells with small crumpled terminals (**Fig. A1D, E**). Their main neurites sometimes overshot and then came back to the layers 2 and 3, forming characteristic loops. Based on their extensive connectivity to LC11 (Keleş et al., 2020; Tanaka and Clark, 2020) and distinctive morphology (Fischbach and Dittrich, 1989) (**Fig. A1E, F**), these clusters most likely correspond to T3. T3 is a cholinergic (Konstantinides et al., 2018) neuron that connects proximal medulla (M9) and distal lobula, is an ON-OFF cell, and is tightly tuned to small moving objects (Keleş et al., 2020; Tanaka and Clark, 2020). T3 neurons receive cholinergic inputs from both ON (Mi1) and OFF (Tm1) cells (Takemura et al., 2015), which likely implement their ON-OFF property.

**Figure A1.**
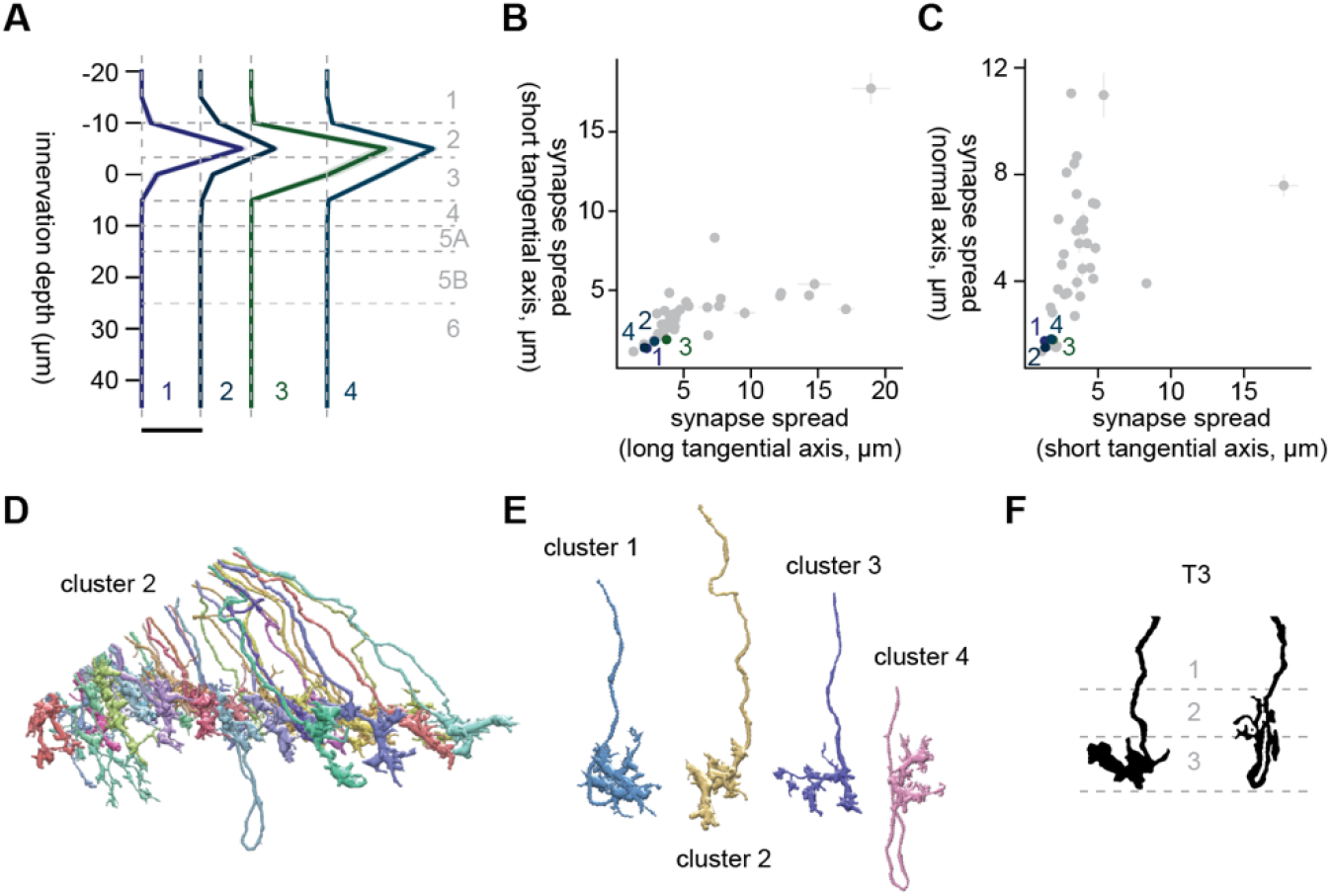
Morphology of clusters 1 through 4, likely corresponding to T3. (A) Mean innervation depth and (B, C) synapse spread along the three axes of clusters 1 through 4. Error bars indicate standard error of the mean. (A) Vertical dotted line indicates zero synapses. The thick horizontal bar indicates 20 synapses. (D) A population of cluster 2 cells, viewed from a tangential direction. (E) Representative examples of cells in clusters 1 through 4. (F) Morphology of T3 axon terminals, from Fischbach & Dittrich (1989).

### Clusters 5 through 7

The branch containing clusters 5 through 15 was characterized by high synapse spread along the long tangential axis, reflecting their elongated morphology (**Fig. A2A**). Clusters 5 through 7 had LC4, a loom sensitive visual projection neuron (Ache et al., 2019; von Reyn et al., 2014) as their main postsynaptic target (11.5, 16.9, and 31.6 synapses/cell for clusters 5, 6, and 7, respectively). Cluster 5 was a homogeneous cluster mostly consisting of monostratified cells with T-shaped, spiny axon terminals extending along the long tangential axis, which innervated slightly shallower than the clusters 1 through 4 (T3) (**Fig. A2A-E**). Its other major targets included LC11 (3.8 synapses/cell), LC18 (3.5 synapses/cell), Li15 (2.6 synapses/cell), and LPLC1 (2.5 synapses/cell). Based on their morphology (Fischbach and Dittrich, 1989) (**Fig. A2E, F**), and connectivity to LC11 (Keleş et al., 2020) cluster 5 most likely represents T2 neurons. T2 neurons, similar to T3 neurons, are cholinergic (Konstantinides et al., 2018) ON-OFF cells with tight size tuning (Keleş et al., 2020; Tanaka and Clark, 2020). T2 neurons, again similar to T3 neurons, have bushy dendrite in the medulla layer 9, but also have additional dendrites in proximal medulla (Fischbach and Dittrich, 1989). Their known inputs include both excitatory ON and OFF cells (Mi1, Tm2, L4, and L5) (Takemura et al., 2015).

Clusters 6 and 7 were somewhat heterogeneous clusters with small membership (**Fig. A2G**). They were slightly more vertically diffuse than cluster 5 (**Fig. A2A, C**). Cluster 6 consisted of at least two cell types: small vertical cells with Y-shaped ending at Lo4 and swelling at Lo2 (**Fig. A2H**), and thin monostratified cells in L4 with long, knobby neurites branching perpendicularly (**Fig. A2J**). Cluster 7 was similar to cluster 6, but contained horizontal spiny cells in Lo2, which appeared to be fragmented parts of larger cells (**Fig. A2K**). Their major targets after LC4 were LPLC1 (2.1 synapses/cell) for cluster 6 and LC12 (2.0 synapses/cell) for cluster 7, but these synapse counts were about an order of magnitude lower than their connectivity to LC4. The terminal morphology of the vertical cells resembled Tm4 (Fischbach and Dittrich, 1989) (**Fig. A2I**), although Tm4 appears to have more extensive swelling in Lo1. TmY2 also had similar Y-shaped ending (**Fig. A2I**), but they have swelling in Lo3 rather than Lo2 (Fischbach and Dittrich, 1989).

**Figure A2.**
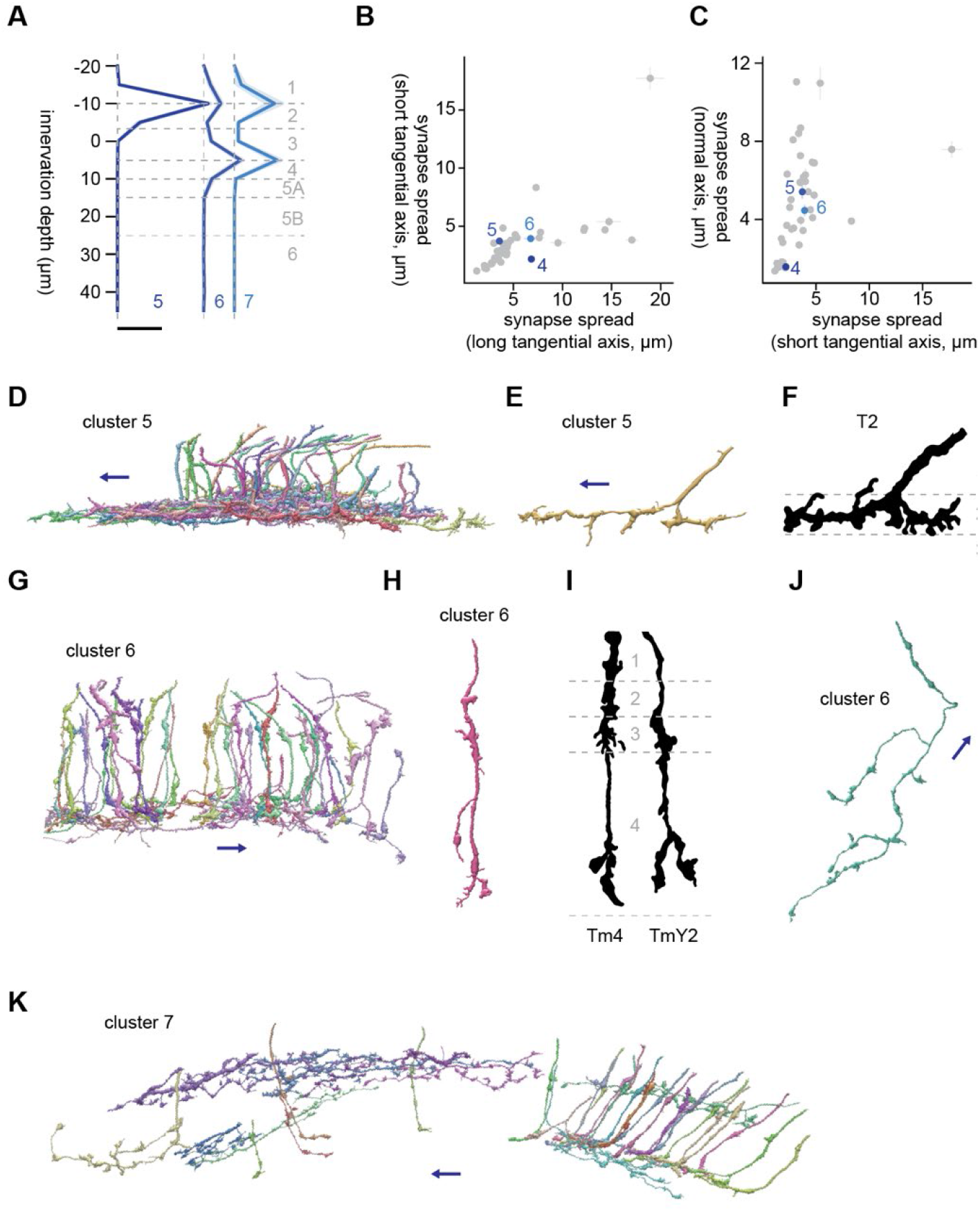
Morphology of clusters 5 through 7. (A) Mean innervation depth and (B, C) synapse spread along the three axes of clusters 5 through 7. Error bars indicate standard error of the mean. (A) Vertical dotted line indicates zero synapses. The thick horizontal bar indicates 20 synapses. (D) A population of cluster 5 cells, viewed from a tangential direction. (E) A representative cluster 5 cell. (F) Morphology of T2 axon terminals, from Fischbach & Dittrich (1989). (G) A population of cluster 6 cells, viewed from a tangential direction. (H) A typical “vertical” cell in cluster 6. (I) Morphology of Tm4 and TmY2, from Fischbach & Dittrich (1989). (J) A typical “monostratified” cell in cluster 6, viewed from a normal direction. (K) A population of cluster 7 cells, viewed from a tangential direction. The blue arrows indicate the ventral direction throughout.

### Clusters 8 through 11

Clusters 8 through 11 were characterized by their large synapse spread along the long tangential axis, as well as the normal axis (**Fig. A3B, C**). They generally innervated deeper than Lo4 (**Fig. A3A**), and shared connectivity to LC10 (7.6, 9.0, 3.0, and 1.4 synapses/cell for clusters 8, 9, 10, and 11, respectively), a group of projection neurons involved in male courtship rituals (Ribeiro et al., 2018; Sten et al., 2021). Cluster 8 consisted of large, tangentially isotropic neurons monostratified in Lo6, likely representing an as yet unlabeled lobula intrinsic (Li) neuron type (**Fig. A3D, E**). Their somata sat on the ventral end of the lobula, and their main neurite entered proximally into lobula (**Fig. A3D**). Each of these cells covered about a quarter of the tangential extent of lobula (**Fig. A3E**). Their major postsynaptic targets after LC10 were LC20 (5.9 synapses/cell), mALC2 (5.2 synapses/cell), LT52 (5.2 synapses/cell), and Li12 (4.4 synpses/cell).

Cluster 9 was a highly anisotropic cluster with synapse spread of 12.2 and 4.8 microns along short and long tangential axes (**Fig. A3B**). The dominant cell type of this cluster had a long, knobby neurites branching at acute angles in Lo5B, which entered lobula ventrally and then bent dorsally (**Fig. A3F, G**). The major postsynaptic targets of this cluster after LC10 were LC13 (5.1 synapses/cell), Li19 (3.8 synapses/cell), LC6 (2.3 synapses/cell), and LT57 (2.0 synapses/cell).

Cluster 10 was another highly anisotropic cluster with synapse spread of 12.2 and 4.6 microns along short and long tangential axes (**Fig. A3B**). The major postsynaptic targets of cluster 10 other than LC10 were Li16 (5.5 synapses/cell), Li12 (4.5 synapses/cell), LC15 (2.7 synapses/cell), and Li13 (2.4 synapses/cell). The dominant cell type of this cluster had a linear neurite with little branching that entered lobula dorsally and then bent ventrally almost perpendicularly (**Fig. A3H**). While this cluster appears to be bistratified in Lo4 and Lo5B as a population (**Fig. A3H**), closer inspection revealed variations in the morphology of single cells (**Fig. A3I**): Some cells were genuinely bistratified, but the branches were often longer in one layer than the other. Some other cells entirely lacked branching. Interestingly, some of these non-branching cells initially entered Lo5B and then switched to Lo4. This long, linear terminal morphology resembled TmY9 (Fischbach and Dittrich, 1989) (**Fig. A3J**). TmY9 is known to have long, linear neurites similar to its lobula terminal in medulla and lobula plate as well (Fischbach and Dittrich, 1989). Another known cell type with long, linear terminal oriented ventrally is Tm28 (Fischbach and Dittrich, 1989) (**Fig. A3J**), although they appear to innervate a little deeper. Note that, although the drawing of TmY9 in Fischabch & Dittrich (1989) lacks branching, *Musca* Y17 cells, a putative TmY9 homologue, has a bistratified branching (Douglass and Strausfeld, 1998), similar to what is shown here.

Cluster 11 was a highly heterogeneous cluster with a variety of cell types in deeper layers of lobula with elongated morphology along the long tangential axis (**Fig. A3K**). It appeared to contain cells similar to those in clusters through 8 to 10. The largest postsynaptic target of this cluster was LC10, but the mean synapse count per cell was only 1.4, likely reflecting the heterogeneity of the cluster.

**Figure A3.**
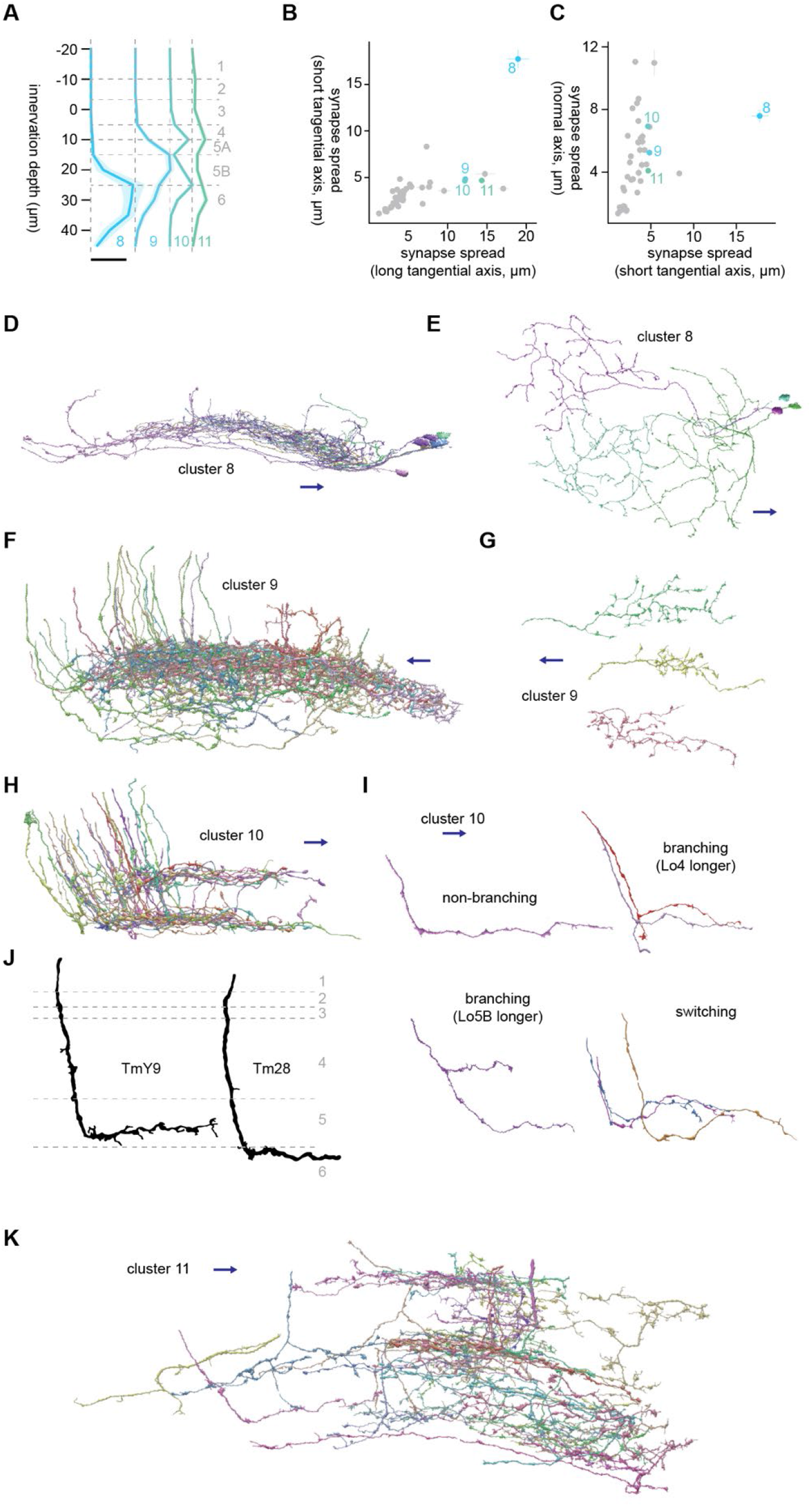
Morphology of clusters 8 through 11. (A) Mean innervation depth and (B, C) synapse spread along the three axes of clusters 8 through 11. Error bars indicate standard error of the mean. (A) Vertical dotted line indicates zero synapses. The thick horizontal bar indicates 20 synapses. (D) A population of cluster 8 cells, viewed from a tangential direction. (E) Representative cluster 8 cells, viewed from a normal direction. (F) A population of cluster 9 cells, viewed from a tangential direction. (G) Representative cluster 9 cells, viewed from a normal direction. (H) A population of cluster 10 cells, viewed from a tangential direction. (I) Representative cluster 10 cells, viewed from a normal direction. Four different subtypes are shown, as labeled. (J) Morphology of TmY9 and Tm28 axon terminals, from Fischbach & Dittrich (1989). (K) A population of cluster 11 cells, viewed from a tangential direction. The blue arrows indicate the ventral direction throughout.

### Clusters 12 through 15

Clusters 12 through 15 are another set of clusters with large synapse spread along the long tangential axis (**Figs. 3A, A4B, C**). These clusters innervated shallower around Lo2 to Lo4 (**Fig. A4A**), unlike the previous 4 clusters. Clusters 12 and 13 were homogeneous clusters dominated by vertically diffuse cells that shared connection to LT87 (10.9 and 19.0 synapses/cell for clusters 12 and 13, respectively), as well as to LC31 (2.9 and 12.2 synapses/cell for clusters 12 and 13, respectively) and LC17 (2.4 and 9.9 synapses/cell for clusters 12 and 13, respectively) (**Fig. A4D**). These cells had a meandering main neurite with many minor branches (**Fig. A4E**). In addition, they had a long secondary branch sticking out ventrally from the main neurite perpendicularly at around Lo3, which was also meandering and with many minor branches (**Fig. A4E**). Among previously documented cells, Tm19 appeared similar to these clusters in that it has an extensive secondary branch (Fischbach and Dittrich, 1989) (**Fig. A4F**), but their innervation appears deeper.

Unlike clusters 12 and 13, clusters 14 and 15 were highly heterogeneous and it was difficult to make out the dominant cell types (**Fig. A4G**). The main postsynaptic targets of these clusters were LPLC1 (19.9 and 5.7 synapses/cell for clusters 14 and 15, respectively). Cluster 14 contained some T2 cells, as can be also seen from the UMAP embedding, where some cluster 14 cells are next to the T2-dominated cluster 5.

**Figure A4.**
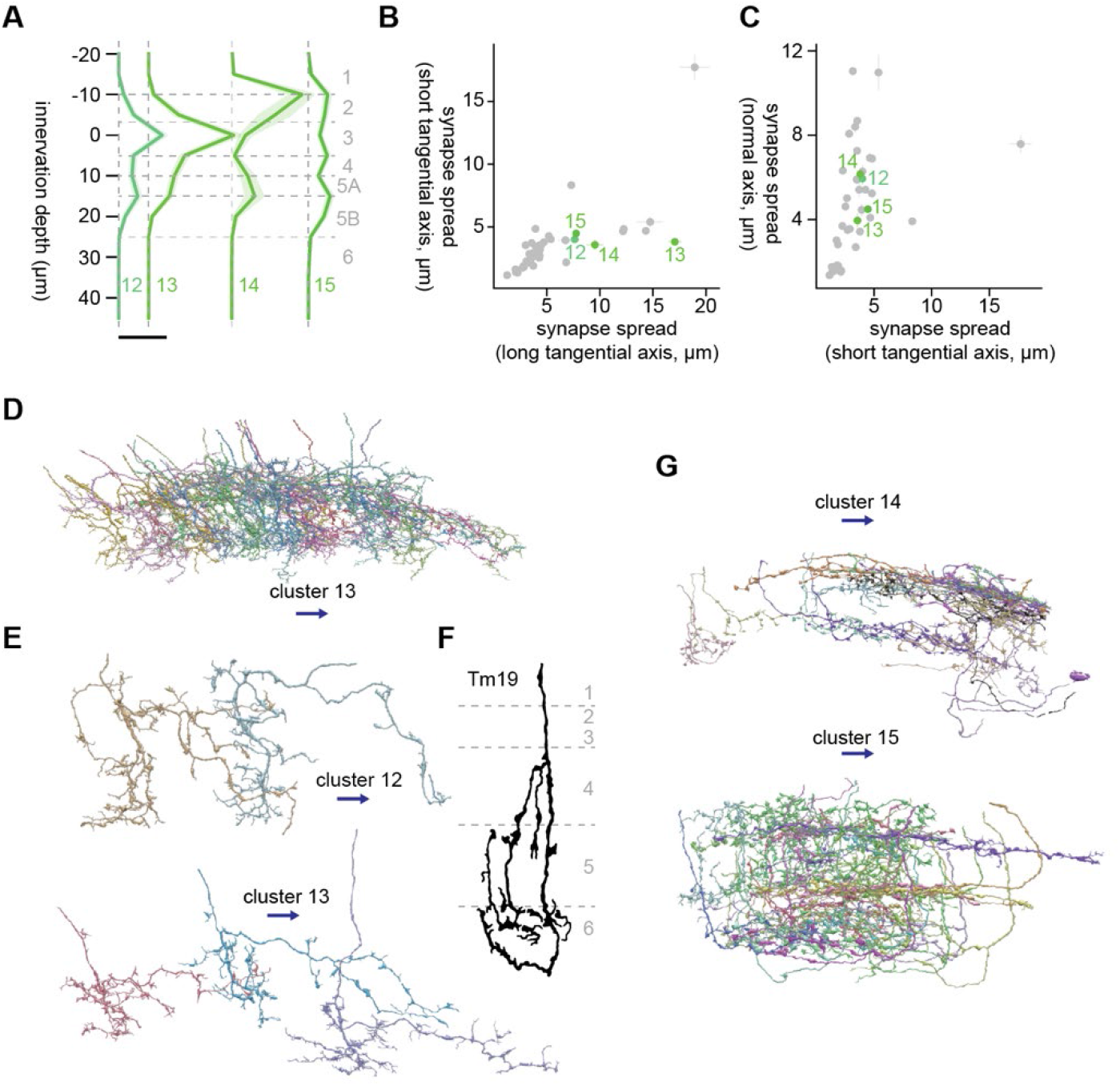
Morphology of clusters 12 through 15. (A) Mean innervation depth and (B, C) synapse spread along the three axes of clusters 12 through 15. Error bars indicate standard error of the mean. (A) Vertical dotted line indicates zero synapses. The thick horizontal bar indicates 20 synapses. (D) A population of cluster 13 cells, viewed from a tangential direction. (E) Representative cluster 12 and 13 cells, viewed from a normal direction. (F) Morphology of Tm19 axon terminals, from Fischbach & Dittrich (1989). (G) A population of cluster 14 and 15 cells, viewed from a tangential direction. The blue arrows indicate the ventral direction throughout.

### Clusters 16 and 17

The branch containing clusters 16 through 22 were characterized by their innervation being relatively restricted to the deep layers, Lo5B and Lo6 (**Fig. A5A**). Clusters 16 and 17 were monostratified clusters in Lo5B with small synapse spread around 3 microns in every direction (**Fig. A5B, C**). They also shared their connectivity to LT58 (8.4 and 12.8 synapses/cell for clusters 16 and 17, respectively), LC10 (4.3 and 4.2 synapses/cell for clusters 16 and 17, respectively), and LC24 (2.6 and 1.8 synapses/cell for clusters 16 and 17, respectively). However, the two clusters appeared to have distinct terminal morphology. Cells in cluster 16 had more tangentially extensive terminals, which consisted of thin, knobby neurites branching perpendicularly, oriented posteriorly (**Fig. A5D, E**). In contrast, the terminals of cluster 17 cells were thicker and branched less (**Fig. A5G, H**). The main neurite of these cells appeared to touch the bottom of Lo5B first and then curl upward, creating a distinctive looped morphology similar to those of T3 (**Fig. A5H**). In terms of connectivity, clusters 17 differed from cluster 16 in that it had a large number of synapses on mALC2 (9.6 synapses/cell). The morphology of cluster 17 cells highly resembled that of Tm20 (Fischbach and Dittrich, 1989) (**Fig. A5I**). The closest match we could find for cluster 16 was Tm11 (**Fig. A5F**), although we are not particularly confident about this identification (Fischbach and Dittrich, 1989). Tm20 is a cholinergic cell (Davis et al., 2020) that is directly postsynaptic to spectrally sensitive photoreceptor R8 (Gao et al., 2008; Takemura et al., 2015), necessary for color learning combinedly with Tm5 subtypes (Melnattur et al., 2014). Other major inputs to Tm20 include L2, L3, C3, Tm1, and Mi4 (Takemura et al., 2015). LT11, a tangential projection neuron necessary for wavelength-specific phototaxis (Otsuna et al., 2014), was not among the strongest postsynaptic target of cluster 17, although LT11 was previously suggested to be downstream of Tm20 based on transsynaptic GFP reconstruction (GRASP) (Lin et al., 2016).

**Figure A5.**
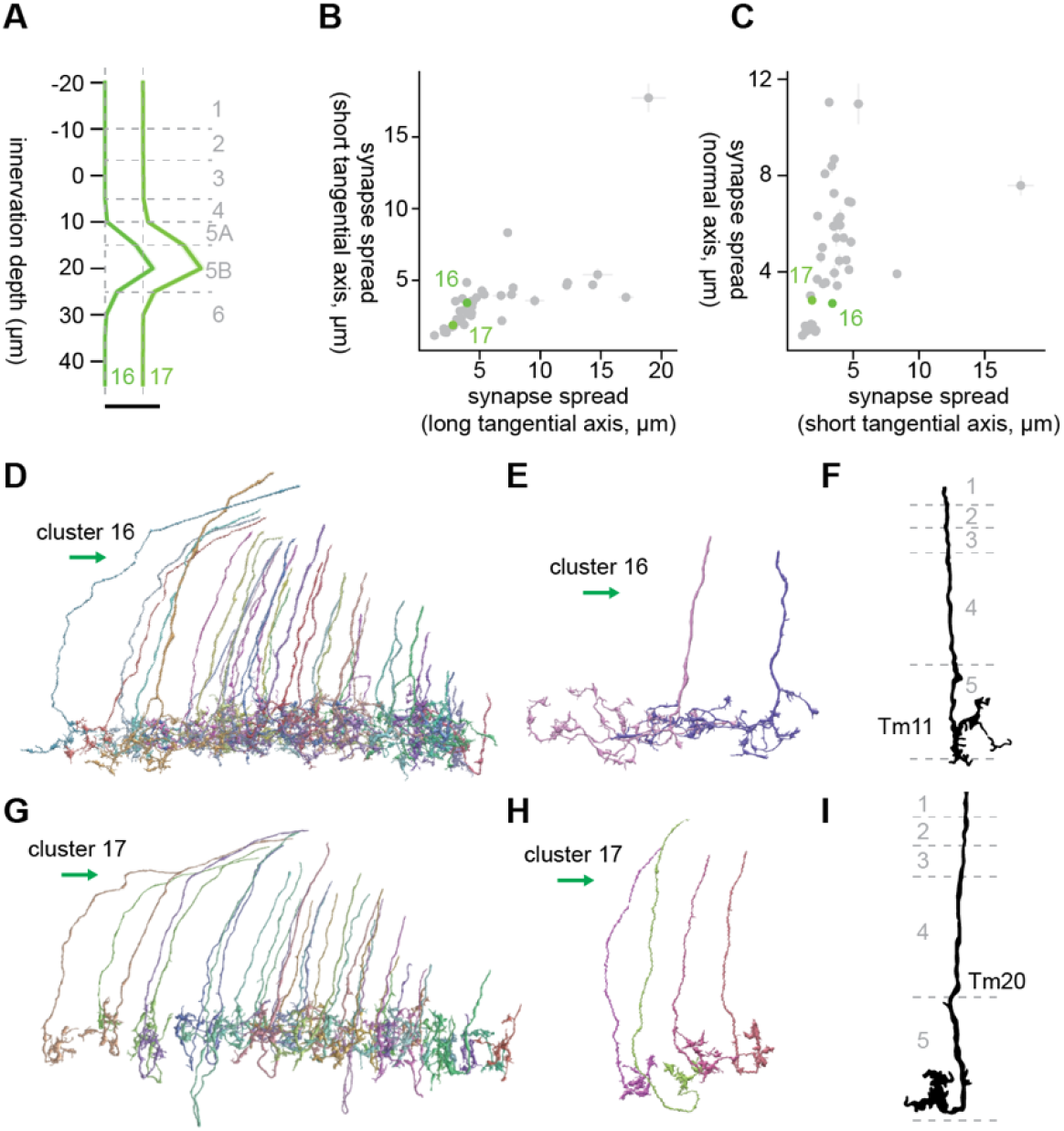
Morphology of clusters 16 and 17. (A) Mean innervation depth and (B, C) synapse spread along the three axes of clusters 16 and 17. Error bars indicate standard error of the mean. (A) Vertical dotted line indicates zero synapses. The thick horizontal bar indicates 20 synapses. (D) A population of cluster 16 cells, viewed from a tangential direction. (E) Representative cluster 16 cells. (F) Morphology of Tm11 axon terminals, from Fischbach & Dittrich (1989). (G) A population of cluster 17 cells, viewed from a tangential direction. (H) Representative cluster 17 cells. (I) Morphology of a Tm20 axon terminal, from Fischbach & Dittrich (1989). The green arrows indicate the anterior direction throughout.

### Clusters 18 and 19

Clusters 18 and 19 were deep monostratified clusters similar to clusters 16 and 17, but they innervated slightly deeper, reaching Lo6 (**Fig. A6A**). They also shared connectivity to mALC2 (14.7 and 7.5 synapses/cell for clusters 18 and 19, respectively), but their postsynaptic targets were distinct beyond that: Cluster 18 synapsed onto LPLC2 (4.8 synapses/cell), LT58 (4.1 synapses/cell), LC22 (3.8 synapses/cell), and LC6 (3.6 synapses/cell), while cluster 19 synapsed onto LC10 (3.6 synapses/cell), mALC1 (2.4 synapses/cell), and Li12 (1.3 synapses/cell). Cluster 18 consisted of homogeneous cells whose terminals had short branches with knobby endings (**Fig. A6D, E**). The dominant cell type of cluster 19 had terminals with fewer branches compared to cluster 18, but they were vertically more diffuse and had small protrusions in shallower layers (**Fig. A6F, G**). The best morphological matches we could find for these clusters were TmY10 (Fischbach and Dittrich, 1989) (**Fig. A6H**), but we are not particularly confident with this identification.

**Figure A6.**
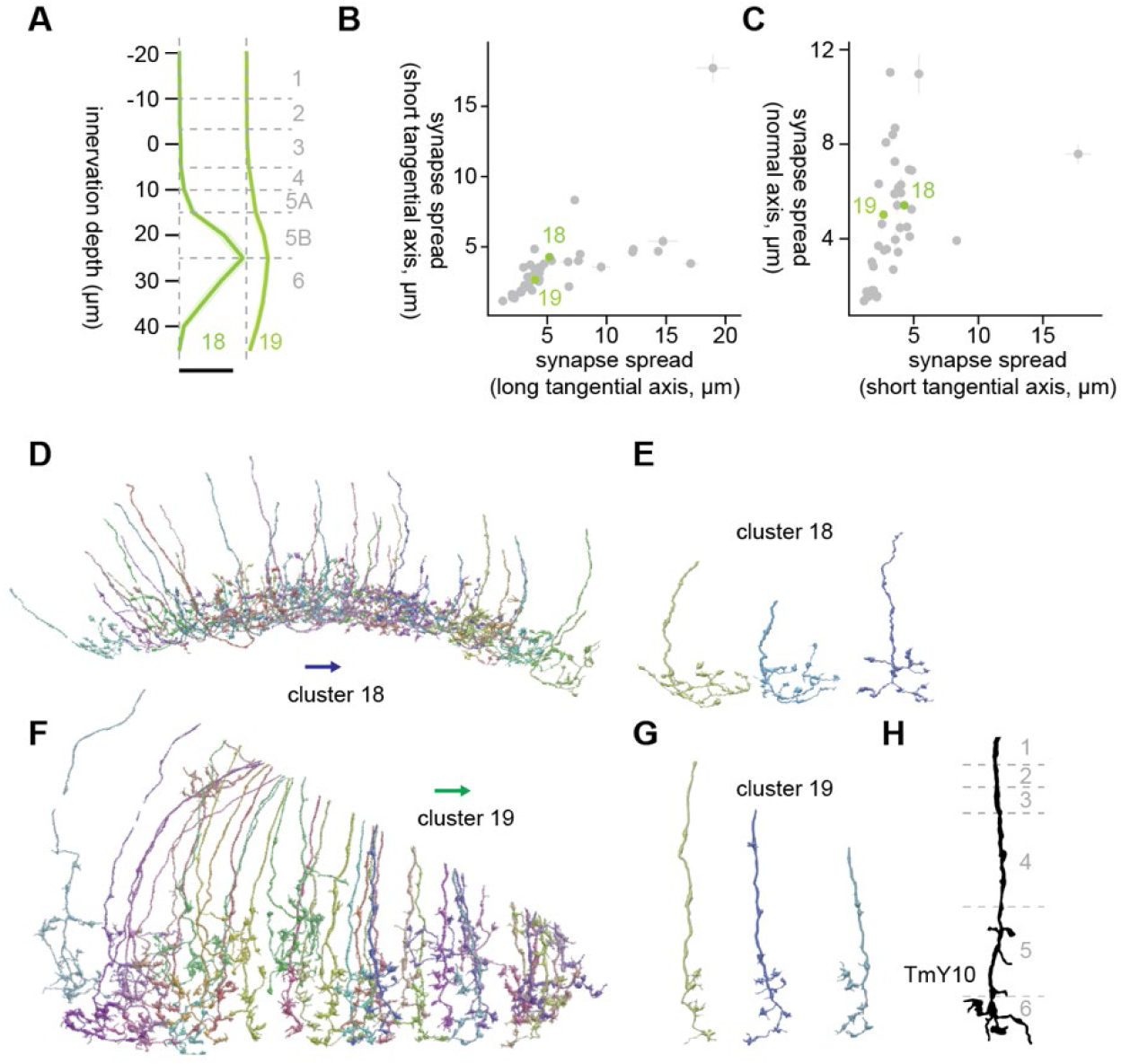
Morphology of clusters 18 and 19. (A) Mean innervation depth and (B, C) synapse spread along the three axes of clusters 18 and 19. Error bars indicate standard error of the mean. (A) Vertical dotted line indicates zero synapses. The thick horizontal bar indicates 20 synapses. (D) A population of cluster 18 cells, viewed from a tangential direction. (E) Representative cluster 18 cells. (F) A population of cluster 19 cells, viewed from a tangential direction. (G) Representative cluster 19 cells. (H) Morphology of a TmY10 axon terminal, from Fischbach & Dittrich (1989). The blue and green arrows respectively indicate the ventral and anterior directions.

### Clusters 20 through 22

Clusters 20 through 22 share their strong connectivity to mALC2 with the previous two clusters (7.5, 24.3, and 19.7 synapses/cell for clusters 20, 21, and 22, respectively). Unlike clusters 16 through 19, they were bistratified (**Fig. A7A**). Cluster 20 consisted of large, lobula intrinsic (Li) neurons bistratified in Lo6 and Lo4/5A, whose cell bodies were on the ventral end of lobula (**Fig. A7D**). Their bistratified morphology, as well as knobby appearance (**Fig. A7D, E**), resembled Li1 (Fischbach and Dittrich, 1989) (**Fig. A7F**). Major postsynaptic targets of this cluster other than mALC2 were LPLC4 (10.7 synapses/cell), LC10 (6.7 synapses/cell), Li13 (5.7 synapses/cell), and LT36 (4.9 synapses/cell).

Cluster 21 consisted of cells with thick neurite, which are bistratified in Lo6 and Lo5A (**Fig. A7A, G, H**). Individual neurons in this cluster had a single side branch in Lo5A and multiple branches in Lo6, all oriented dorsally (**Fig. A7H**). These oriented branches resembled those of Tm8 (Fischbach and Dittrich, 1989) (**Fig. A7I**). The major postsynaptic targets of this cluster after mALC2 were LC10 (15.5 synapses/cell), LT57 (5.1 synapses/cell), LC13 (4.9 synapses/cell), and LT52 (3.7 synapses/cell). Cluster 22 consisted of at least two distinct cell types (**Fig. A7J-L**). One was thick, bistratified cells similar to cluster 21 (**Fig. A7K**), but their branches were not oriented like cluster 21 cells. The other was thin cells that had terminals with branches with knobby endings and a small protrusion at a shallower layer (**Fig. A7L**). Their major postsynaptic targets after mALC2 were LC10 (8.2 synapses/cell), LT52 (4.1 synapses/cell), LT63 (2.6 synapses/cell), and LT51 (2.6 synapses/cell).

**Figure A7.**
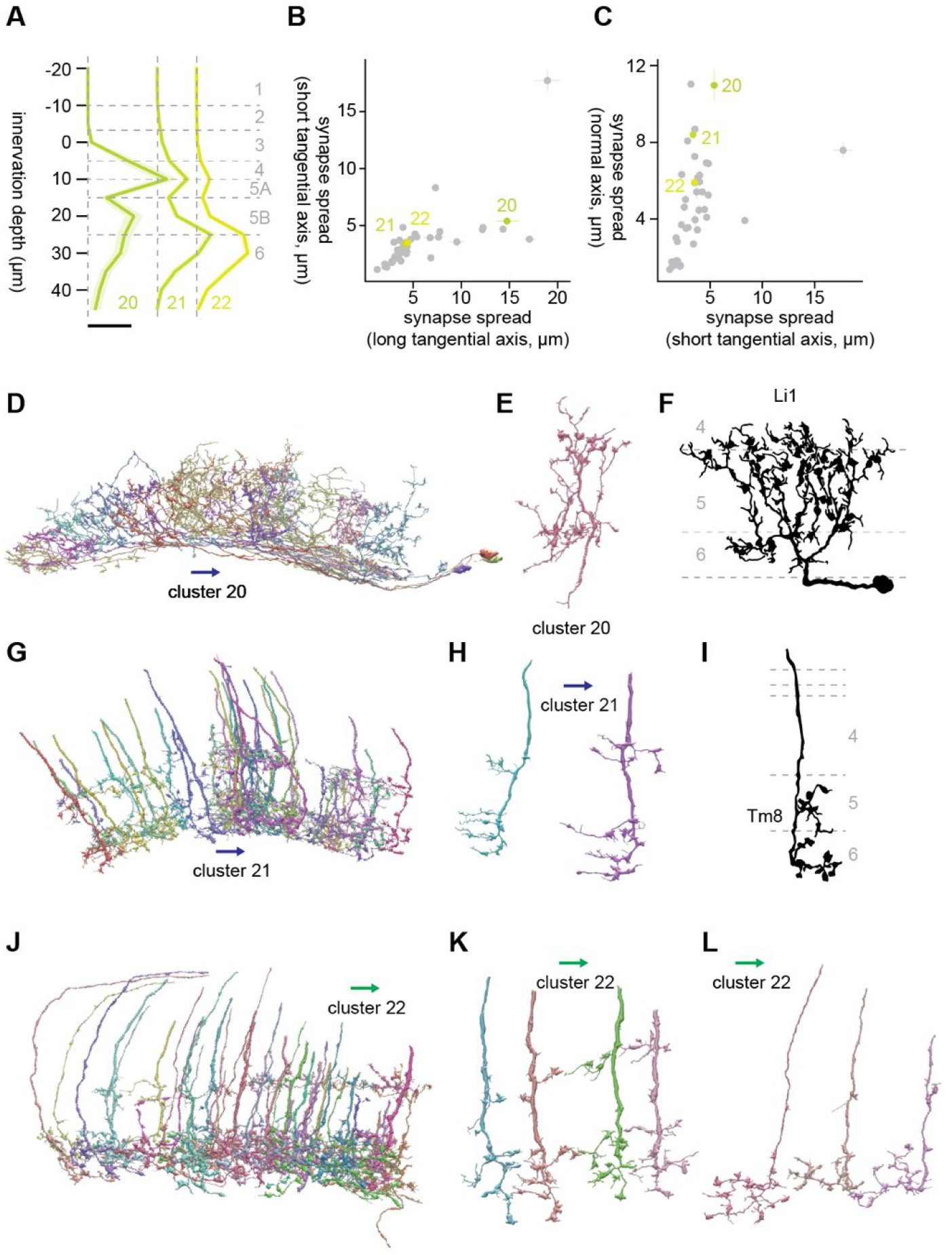
Morphology of clusters 20 through 22. (A) Mean innervation depth and (B, C) synapse spread along the three axes of clusters 20 through 22. Error bars indicate standard error of the mean. (A) Vertical dotted line indicates zero synapses. The thick horizontal bar indicates 20 synapses. (D) A population of cluster 20 cells, viewed from a tangential direction. (E) A representative cluster 20 cell. (F) Morphology of a Li1 cell, from Fischbach & Dittrich (1989). (G) A population of cluster 21 cells, viewed from a tangential direction. (H) Representative cluster 21 cells. (I) Morphology of a Tm8 axon terminal, from Fischbach & Dittrich (1989). (J) A population of cluster 22 cells, viewed from a tangential direction. (K, L) Representative (K) thick and (L) thin cells in cluster 22. The blue and green arrows respectively indicate the ventral and anterior directions.

### Cluster 23

Cluster 23 was highly distinct from the neighboring clusters, as can be seen both from the dendrogram (**Fig 3A**) and UMAP embedding (**Fig. 3E**). Cluster 23 consists of a small monostratified cells in Lo3, with synapse spread of around 2 microns in every direction (**Fig. A8A-C**). Its main postsynaptic targets were Li11 (8.5 synapses/cell), LT1 (4.8 synapses/cell), LC11 (2.6 synapses/cell), LPLC1 (2.1 synapses/cell), and LC21 (1.8 synapses/cell). Visually, cluster 23 appeared to consist of highly homogenous cells with a knobby terminal and a small protrusion at a shallower location (**Fig. A8D, E**). The best morphological match we could find was T2a, a cell type morphologically and transcriptomically related to T2 and T3 (Fischbach and Dittrich, 1989; Keleş et al., 2020; Özel et al., 2020) (**Fig. A8F**). Indeed, this cluster shared its connectivity to LT1, LC11, and LPLC1 with T3, indicating that it could be functionally similar to T3. While functional property of T2a has not been studied, it shares inputs from Mi1 and Tm1 with T3, implying it is also an ON-OFF cell (Takemura et al., 2015). T2a lacks inputs from L cells unlike T2, despite its proximal medulla innervation (Takemura et al., 2015). Another cell type with similar terminal morphology was Tm21 (Fischbach and Dittrich, 1989) (**Fig. A8F**).

**Figure A8.**
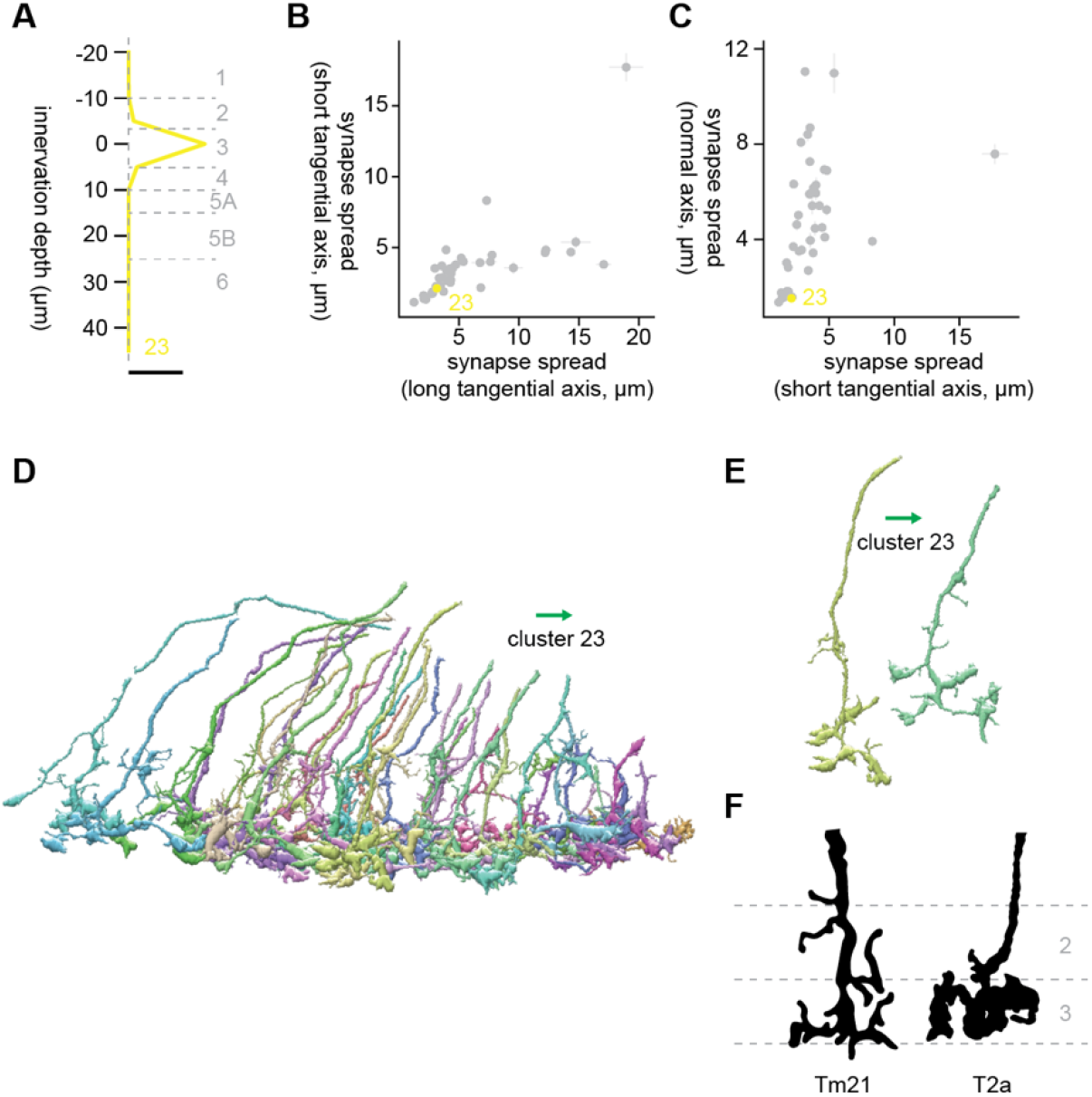
Morphology of cluster 23. (A) Mean innervation depth and (B, C) synapse spread along the three axes of cluster 23. Error bars indicate standard error of the mean. (A) Vertical dotted line indicates zero synapses. The thick horizontal bar indicates 20 synapses. (D) A population of cluster 23 cells, viewed from a tangential direction. (E) Representative cluster 23 cells. (F) Morphology of Tm21 and T2a cells, from Fischbach & Dittrich (1989). The green arrows indicate the anterior direction.

### Clusters 24 and 25

Clusters 24 and 25 were highly heterogeneous clusters, and it was difficult to make out the dominant cell types (**Fig. A9D, F**). The low synapse counts to even their largest targets (cluster 24: 2.16 synapses/cell to mALC2; cluster 25: 1.2 synapses/cell to LC4) and the very diffuse innervation depth profile (**Fig. 12A**) likely reflect this heterogeneity. Of note, cluster 24 contained a set of neurites that entered lobula from the dorsal proximal end as if hugging the bottom of Lo6 (**Fig. A9D, E**). Cluster 25 contained a large number of fragmented neurites around the dent in lobula (**Fig. A9F**) which was likely caused by how it was mounted for scanning. Thus, this cluster may be driven by the imaging methodology rather than by the biology.

**Figure A9.**
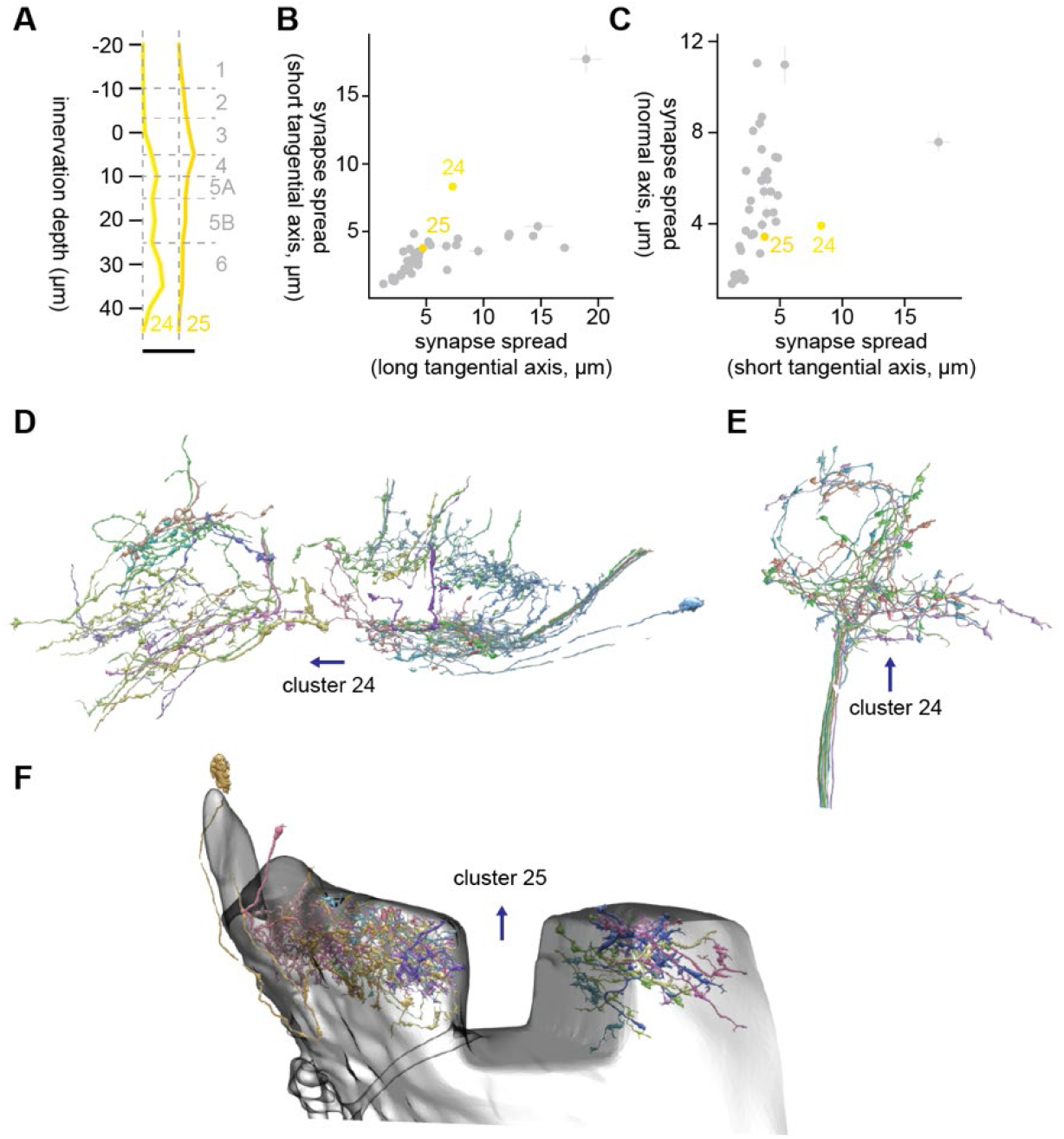
Morphology of clusters 24 and 25. (A) Mean innervation depth and (B, C) synapse spread along the three axes of clusters 24 and 25. Error bars indicate standard error of the mean. (A) Vertical dotted line indicates zero synapses. The thick horizontal bar indicates 20 synapses. (D) A population of cluster 24 cells, viewed from a tangential direction. (E) Cluster 24 cells whose process entered lobula from proximal and dorsal end. (F) A population of cluster 25 cells around the dent in lobula. The transparent shell represents the outer boundary of lobula. The blue arrows indicate the ventral direction.

### Clusters 26 through 29

Clusters 26 through 29 were deep monostratified clusters in Lo5A/B (**Fig. A10A**), similar to clusters 16 through 19 (**Figs. A5, 6**). There terminals were relatively compact, with synapse spread around 3 microns in every direction (**Fig. A10B, C**). The major postsynaptic targets of cluster 26 were Li19 (7.2 synapses/cell), LC17 (4.4 synapses/cell), LT79 (2.7 synapses/cell), LPLC2 (2.5 synapses/cell), and LC6 (1.6 synapses/cell). The dominant cell type of this cluster had thick neurites with small protrusions and terminals oriented ventrally with a couple of branching (**Fig. A10D, E**). These morphological features resembled those of Tm5Y (Fischbach and Dittrich, 1989) (**Fig. A10F**).

Cluster 27 was somewhat heterogeneous (**Fig. A10G**), and the only major postsynaptic target with more than single synapse/cell was LC10 (9.6 synapses/cell). The dominant cell type had thin processes and terminals whose short branches had knobby endings (**Fig. A10H**). These features were similar to the thin cells in cluster 22 (**Fig. A7K**), but cluster 27 cells were less tangentially extensive and positioned slightly shallower. Cluster 28 was also dominated by thin, monostratified cells in Lo5B (**Fig. A10I, J**), which had knobby endings, similar to cluster 27. However, postsynaptic targets of cluster 28 were distinct from cluster 27: its major targets were LC6 (6.3 synapses/cell), LPLC2 (2.3 synapses/cell), LC10 (2.1 synapses/cell), Li19 (1.7 synapses/cell), and LC16 (1.5 synapses/cell). Cluster 29 was a large, heterogeneous cluster with deep monostratified neurons (**Fig. A10K**). It contained cells similar to cluster 27 (**Fig. A10H, M)**, as well as to cluster 17 (Tm20) (**Fig. A10L**). LC10 (2.1 synapses/cell) and LC16 (1.3 synapses/cell) were among their major targets. We could not find good morphological match for these monostratified Lo5B cells in Fischbach & Dittrich (1989).

**Figure A10.**
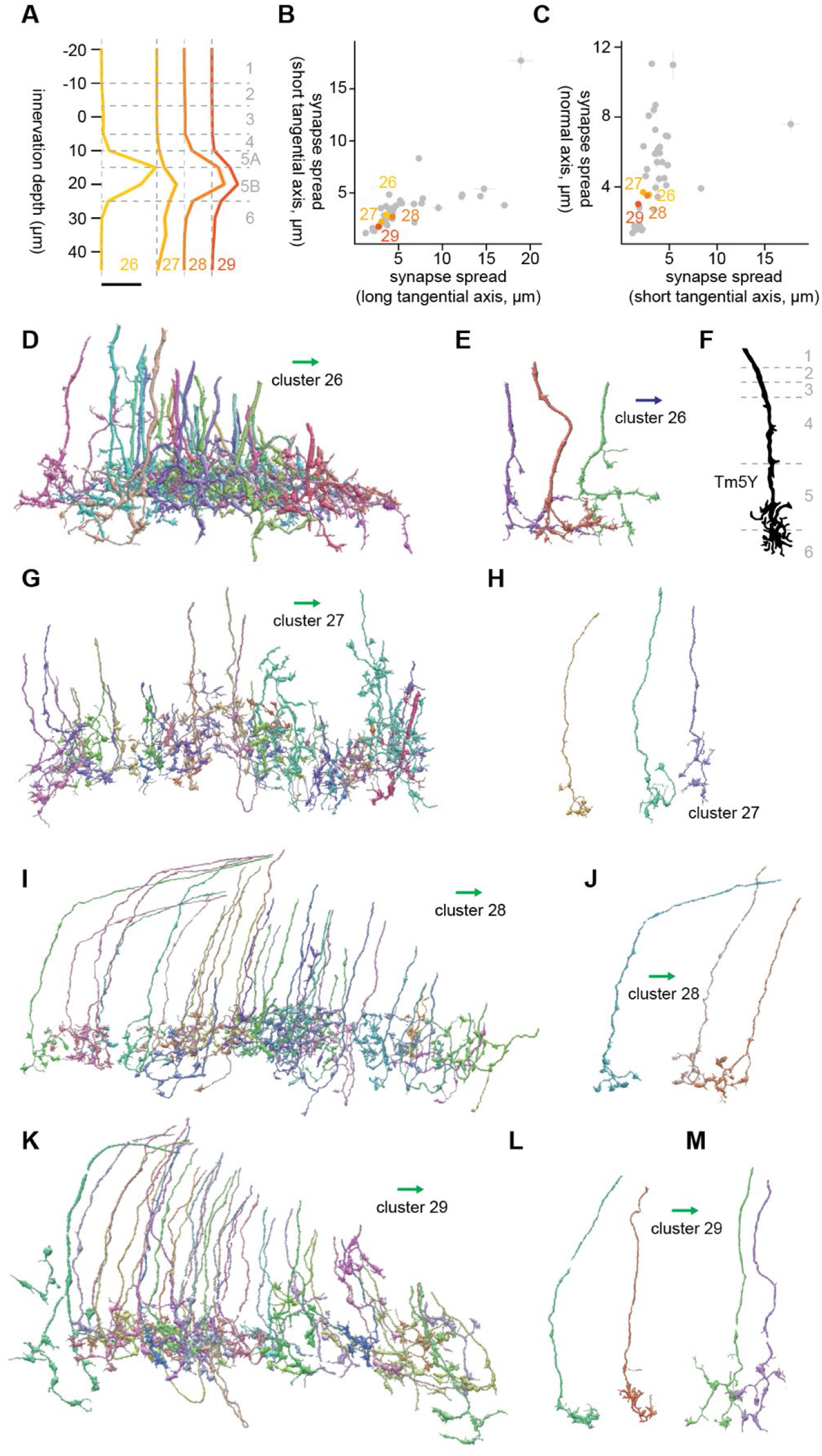
Morphology of clusters 26 through 29. (A) Mean innervation depth and (B, C) synapse spread along the three axes of clusters 26 through 29. Error bars indicate standard error of the mean. (A) Vertical dotted line indicates zero synapses. The thick horizontal bar indicates 20 synapses. (D) A population of cluster 26 cells, viewed from a tangential direction. (E) A representative cluster 26 cell. (F) Morphology of a Tm5Y cell, from Fischbach & Dittrich (1989). (G) A population of cluster 27 cells, viewed from a tangential direction. (H) Representative cluster 27 cells. (I) A population of cluster 28 cells, viewed from a tangential direction. (J) Representative cluster 28 cells. (K) A population of cluster 29 cells, viewed from a tangential direction. (L, M) Representative examples of two cell types included in cluster 29. (L) These cells were similar to cluster 16 cells (Tm20). The blue and green arrows respectively indicate the ventral and anterior directions.

### Clusters 30 and 31

Clusters 30 and 31 were only clusters with substantial Lo1 innervation (**Fig. A11A**). Lo1 appears to be a special layer dedicated to OFF edge motion detection, housing T5 and its inputs but avoided by other cell types, including lobula VPNs (Fischbach and Dittrich, 1989; Shinomiya et al., 2019, 2014; Wu et al., 2016). This is similar to how M10, the putative evolutionary precedent of Lo1 (Shinomiya et al., 2015), is dedicated to T4 and their inputs but avoided by other cells (Fischbach and Dittrich, 1989). Cluster 30 was a highly homogeneous cluster of small monostratified terminals (**Fig. A11D**), whose only major postsynaptic target was CT1 (20.9 synapses/cell). CT1 receives inputs from Tm1 as well as Tm9 in lobula (Shinomiya et al., 2019). The terminal morphology of cluster 30 cells resembled Tm9 more than Tm1 (Fischbach and Dittrich, 1989) (**Fig. A11E**).

Cluster 31 was a large heterogeneous cluster mostly confined to Lo1 (**Fig. A11A, F**). This cluster appears as multiple distant clusters on the UMAP embedding (**Fig. 3E**), also likely reflecting its heterogeneity. Upon visual inspection, we found oriented neurites with dense branching similar to T5 dendrites, as well as small terminals that resembled Tm1, 2, and 9 (**Fig. A11G**). On average, cluster 31 did not have more than single synpase/cell to any of the labeled lobula neurons included in the current analysis. This observation supports the idea that Lo1 is the dedicated layer for motion detection by T5 and is relatively independent from the rest of lobula.

**Figure A11.**
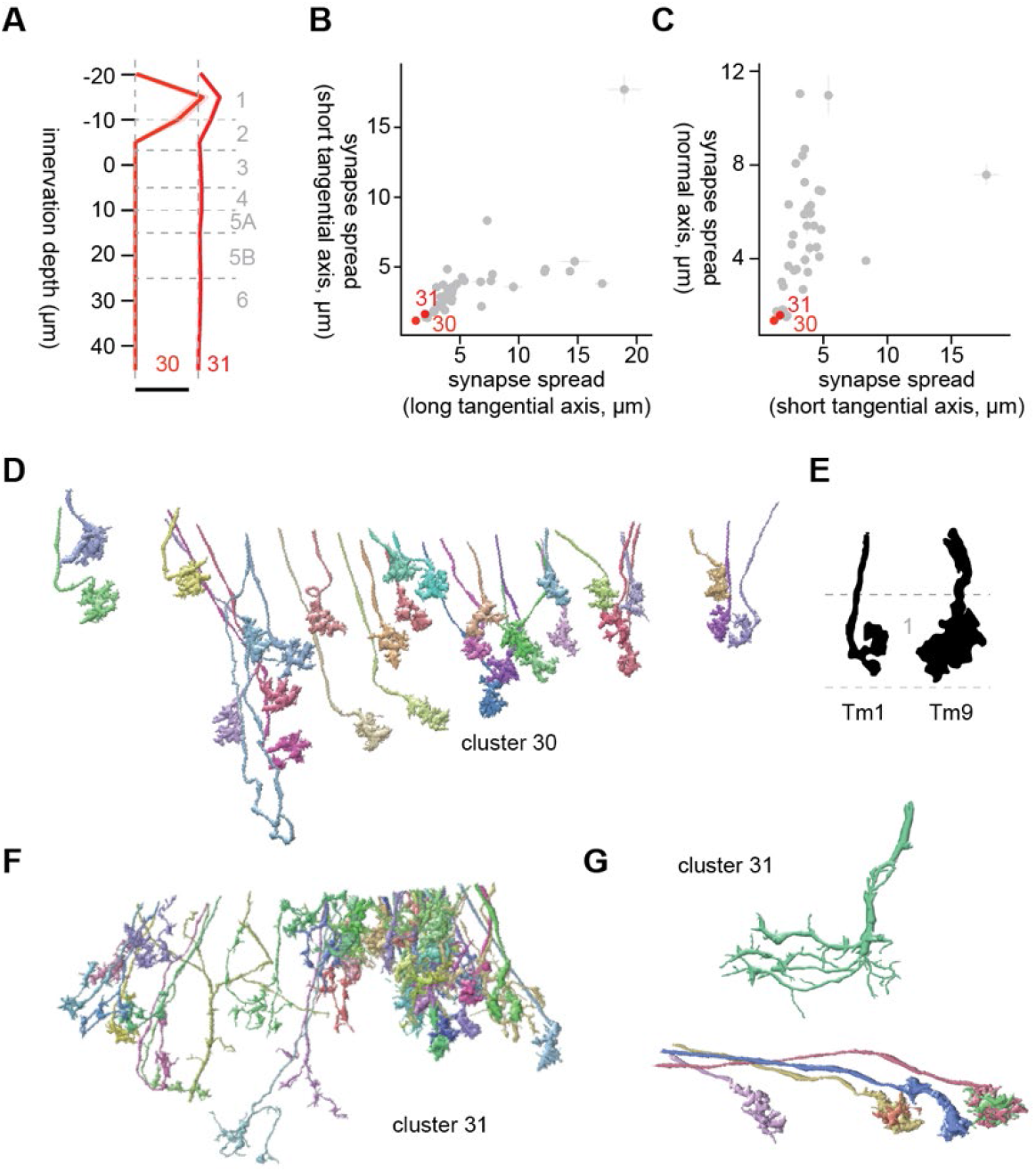
Morphology of clusters 30 and 31. (A) Mean innervation depth and (B, C) synapse spread along the three axes of clusters 30 and 31. Error bars indicate standard error of the mean. (A) Vertical dotted line indicates zero synapses. The thick horizontal bar indicates 20 synapses. (D) The entire population of cluster 30 cells. (E) Morphology of Tm1 and Tm9 axon terminals, from Fischbach & Dittrich (1989). (F) A population of cluster 31 cells. (G) Examples of cluster 31 cells that resembled T5 (*top*) and their inputs (*bottom*).

### Cluster 32 through 34

Clusters 32 through 40 were characterized by their tangentially compact and vertically diffuse morphology (**Fig. A12B, C**). The axon terminals of clusters 32 through 34 were in Lo5A/B, but they also had synapses in shallower layers (**Fig. A12A**). The major postsynaptic targets of cluster 32 were Li19 (12.9 synapses/cell), LC17 (9.3 synapses/cell), LT79 (8.4 synapses/cell), LPLC2 (8.1 synapses/cell), and LPLC1 (7.4 synapses/cell). These cell types overlapped with the main synaptic targets of cluster 26. In fact, the morphology of cluster 32 cells, with its thick neurites with small protrusions and somewhat densely branching terminals (**Fig. A12D, E**), resembled that of cluster 26 cells (**Fig. A10D, E**). Continuity of clusters 26 and 32 can also be seen from the UMAP embedding (**Fig. 3E**). The best morphological match we could find for these clusters was Tm5Y (Fischbach and Dittrich, 1989) (**Fig. A12F**).

Clusters 33 and 34 shared their connectivity to LT11 (20.7 and 11.2 synapses/cell for clusters 33 and 34, respectively), LC25 (7.2 and 4.2 synapses/cell for clusters 33 and 34, respectively), LC11 (5.8 and 1.4 synapses/cell for clusters 33 and 34, respectively), LC15 (2.3 and 1.1 synapses/cell for clusters 33 and 34, respectively), LC26 (1.6 and 1.1 synapses/cell for clusters 33 and 34, respectively). These were the only clusters with strong LT11 connectivity, which most likely made these clusters highly distinct from everything else on the UMAP embedding (**Fig. 3E**). Both clusters were visually homogeneous, consisting of cells characterized by modestly branching terminals in Lo5A/B and straight main neurite with small protrusions in shallower layers (**Fig. A12G-I**). Given their extensive connectivity to LT11, these clusters likely represent Tm5 neurons (Lin et al., 2016). Tm5 neurons, consisting of three subtypes Tm5a/b/c, are glutamatergic neurons postsynaptic to spectrally sensitive photoreceptors R7 and R8 (Gao et al., 2008). Tm5 subtypes, alongside with Tm20, have been shown to be combinedly necessary for color learning in flies (Karuppudurai et al., 2014; Melnattur et al., 2014). Among these subtypes, Tm5c, resembled the cluster 33 and 34 cells the most in that it had a small protrusion in a shallower layer in addition to Lo5 (Gao et al., 2008).

**Figure A12.**
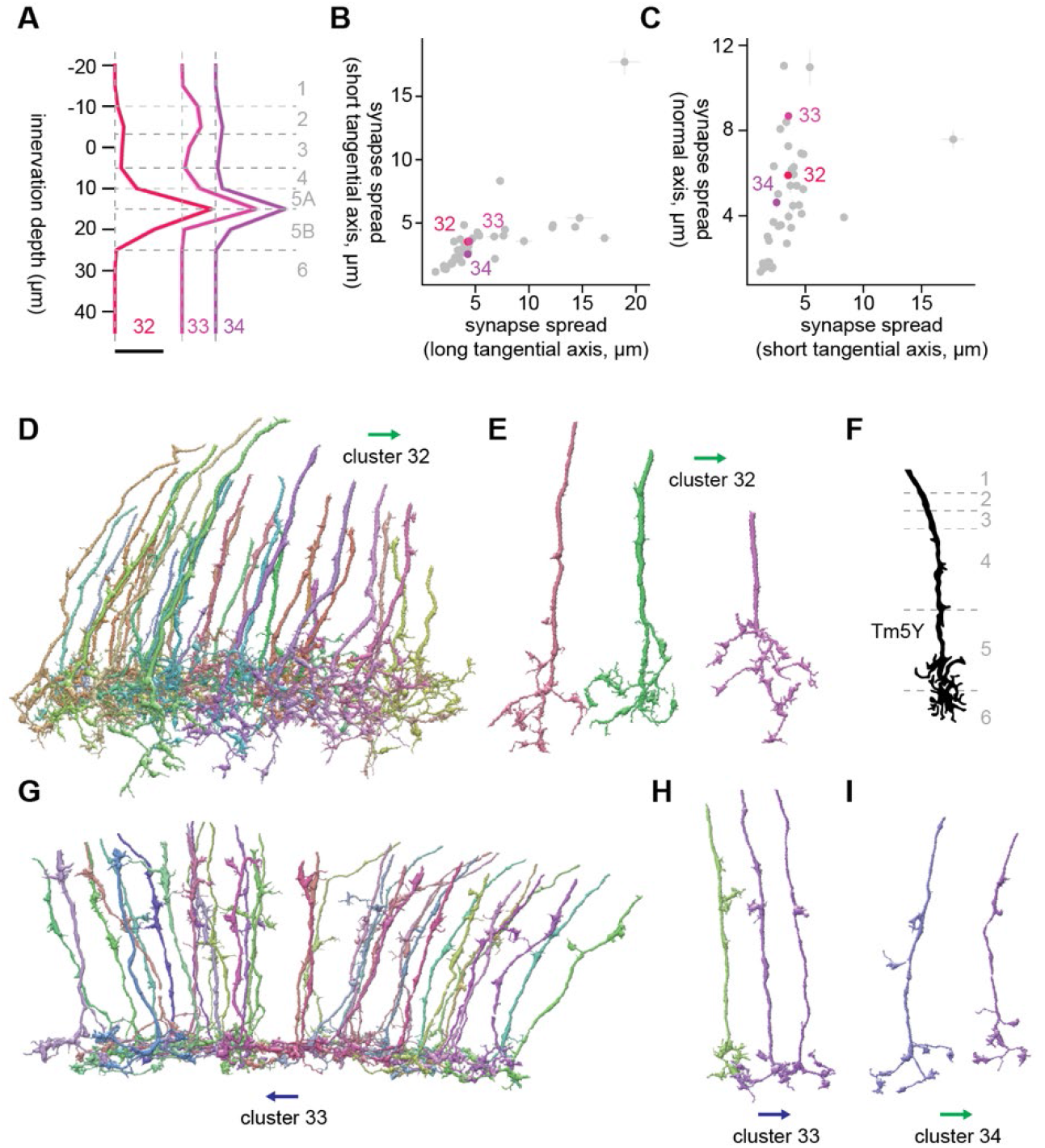
Morphology of clusters 32 through 34. (A) Mean innervation depth and (B, C) synapse spread along the three axes of clusters 32 through 34. Error bars indicate standard error of the mean. (A) Vertical dotted line indicates zero synapses. The thick horizontal bar indicates 20 synapses. (D) A population of cluster 32 cells. (E) Representative examples of cluster 32 cells. (F) Morphology of a Tm5Y axon terminal, from Fischbach & Dittrich (1989). (G) A population of cluster 32 cells. (H, I) Examples of (H) cluster 33 and (I) cluster 34 cells. The blue and green arrows respectively indicate the ventral and anterior directions.

### Clusters 35 through 37

Clusters 35 through 37 were another set of vertically diffuse clusters (**Fig. A13A**), which on average appeared to be bistratified in Lo4/5A and Lo5B/6, but the pattern was less obvious compared to other bistratified clusters. The major postsynaptic targets of cluster 35 were Li16 (7.0 synapses/cell), Li12 (6.1 synapses/cell), LC15 (2.8 synapses/cell), LC10 (1.3 synapses/cell), and Li19 (1.1 synapses/cell). The dominant cell type of cluster 35 had branches sticking out of the main neurite perpendicularly, which aligned with the short tangential axis (**Fig.A13D, E**). This morphology strongly resembled that of TmY11 (Fischbach and Dittrich, 1989) (**Fig. A13F**). The major postsynaptic targets of cluster 36 were LC20 (6.3 synapses/cell), Li12 (2.3 synapses/cell), LC22 (1.2 synapses/cell), LC10 (1.2 synapses/cell), and Li19 (1.0 synapses/cell). The dominant cell type of this cluster appeared more vertically diffuse than cluster 35 (**Fig. A13G, H**), with many short protrusions along its main neurite ranging from L4 to L6. The best morphological match we could find for this cluster was TmY5 (Fischbach and Dittrich, 1989) (**Fig. A13I**). Cluster 37 was another large, heterogeneous cluster consisting of a variety of vertically diffuse neurons resembling cells from clusters 32 through 36 (**Fig. A13J, K**). Similar to cluster 25, cluster 37 contained many fragments around the dent in the sample. The postsynaptic cell types with more than single synapse per cell were LC10 (1.8 synapses/cell) and LC20 (1.4 synapses/cell).

**Figure A13.**
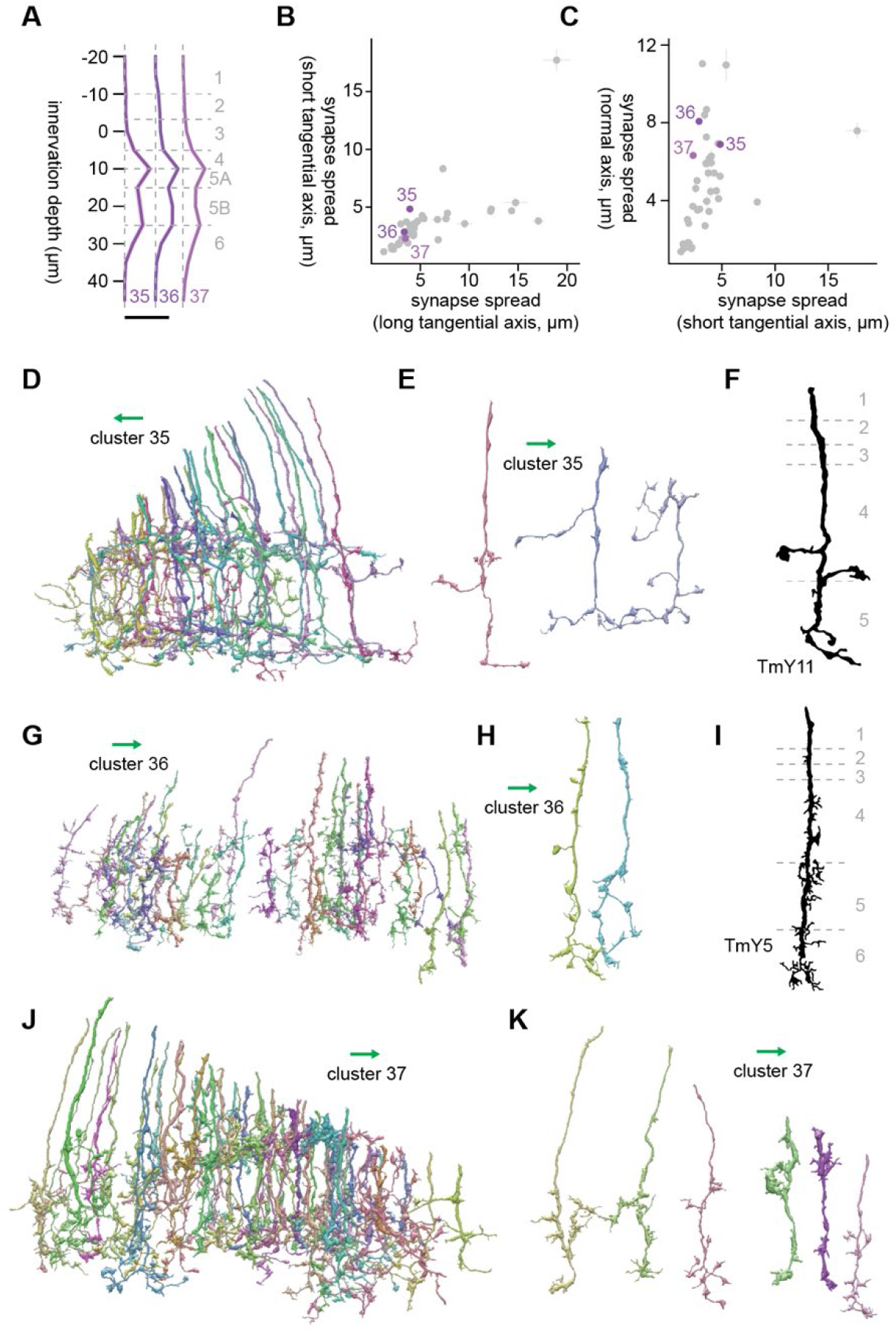
Morphology of clusters 35 through 37. (A) Mean innervation depth and (B, C) synapse spread along the three axes of clusters 35 through 37. Error bars indicate standard error of the mean. (A) Vertical dotted line indicates zero synapses. The thick horizontal bar indicates 20 synapses. (D) A population of cluster 35 cells. (E) Representative examples of cluster 35 cells. (F) Morphology of a TmY11 axon terminal, from Fischbach & Dittrich (1989). (G) A population of cluster 36 cells. (H) Representative examples of cluster 36 cells. (I) Morphology of a TmY5 axon terminal, from Fischbach & Dittrich (1989). (J) A population of cluster 37 cells. (K) Examples cluster 37 cells. The green arrows indicate the anterior direction.

### Clusters 38 through 40

Clusters 38 through 40 were vertically diffuse clusters with shallower layer innervations (**Fig. A14A**). Cluster 38 mainly innervated Lo1/2 and Lo3/4 (**Fig. A14A**). Their major postsynaptic targets were LC4 (4.8 synapses/cell), Li17 (3.9 synapses/cell), mALC1 (3.4 synapses/cell), LPLC1 (1.9 synapses/cell), and LC12 (1.6 synapses/cell). Morphologically, cluster 38 appeared to contain at least two dominant cell types, which is likely why this cluster showed up as two distant clusters on the UMAP embedding (**Fig. 3E**). The first cell type was a short, vertical cell with Y-shaped ending at L4 and swelling in a shallower layer (**Fig. A14D, E**). This morphology, as well as its connectivity, made it resemble cells in cluster 6, which we guessed to be either Tm4 or TmY2 (**Fig. A2G-I**). The continuity of clusters 6 and 38 can be also seen from the UMAP embedding (**Fig. 3E**). The other cell type in cluster 38 reached a deeper layer, likely Lo5A (**Fig. A2F**), and had a knobby appearance with more branching than the first cell type. The best morphological match we could find for this cell type were TmY7 (Fischbach and Dittrich, 1989) (**Fig. A14G**).

Cluster 39 was a small cluster of homogenous cells innervating only the ventral rim of lobula (**Fig. A14H**). These cells had branching terminals with knobby endings (**Fig. A14I**). The major postsynaptic targets of this cluster were Li14 (13.1 synapses/cell), LC10 (7.3 synapses/cell), PS179 (3.9 synapses/cell), LPLC2 (3.3 synapses/cell), and LC14 (2.7 synapses/cell). Extensive connectivity to Li14 as well as PS179 made this cluster unique. It is unclear whether this cluster constitutes a retinotopically specialized cell type by itself. Alternatively, this could be a subset of a cell type whose rest is lost in large clusters like cluster 37, clustered separately due to errors in innervation depth estimation caused by high curvature of lobula layers at the rim.

Cluster 40 was the most vertically diffuse cluster with the synapse spread of 11.0 micron along the normal axis (**Fig. A14C**). This cluster was a homogeneous cluster of bistratified cells in Lo3/4 and Lo6, with minor protrusions in the layers in between (**Fig. A14A, J, K**). The major postsynaptic targets of this cluster were LC10 (4.3 synapses/cell), Li11 (3.1 synapses/cell), LPLC2 (2.2 synapses/cell), LT51 (2.2 synapses/cell), and LC4 (1.8 synapses/cell). The morphology of these cells resembled that of TmY5a (Davis et al., 2020; Fischbach and Dittrich, 1989) (**Fig. A14K, L**), for example, although we are not particularly confident about this identification.

**Figure A14.**
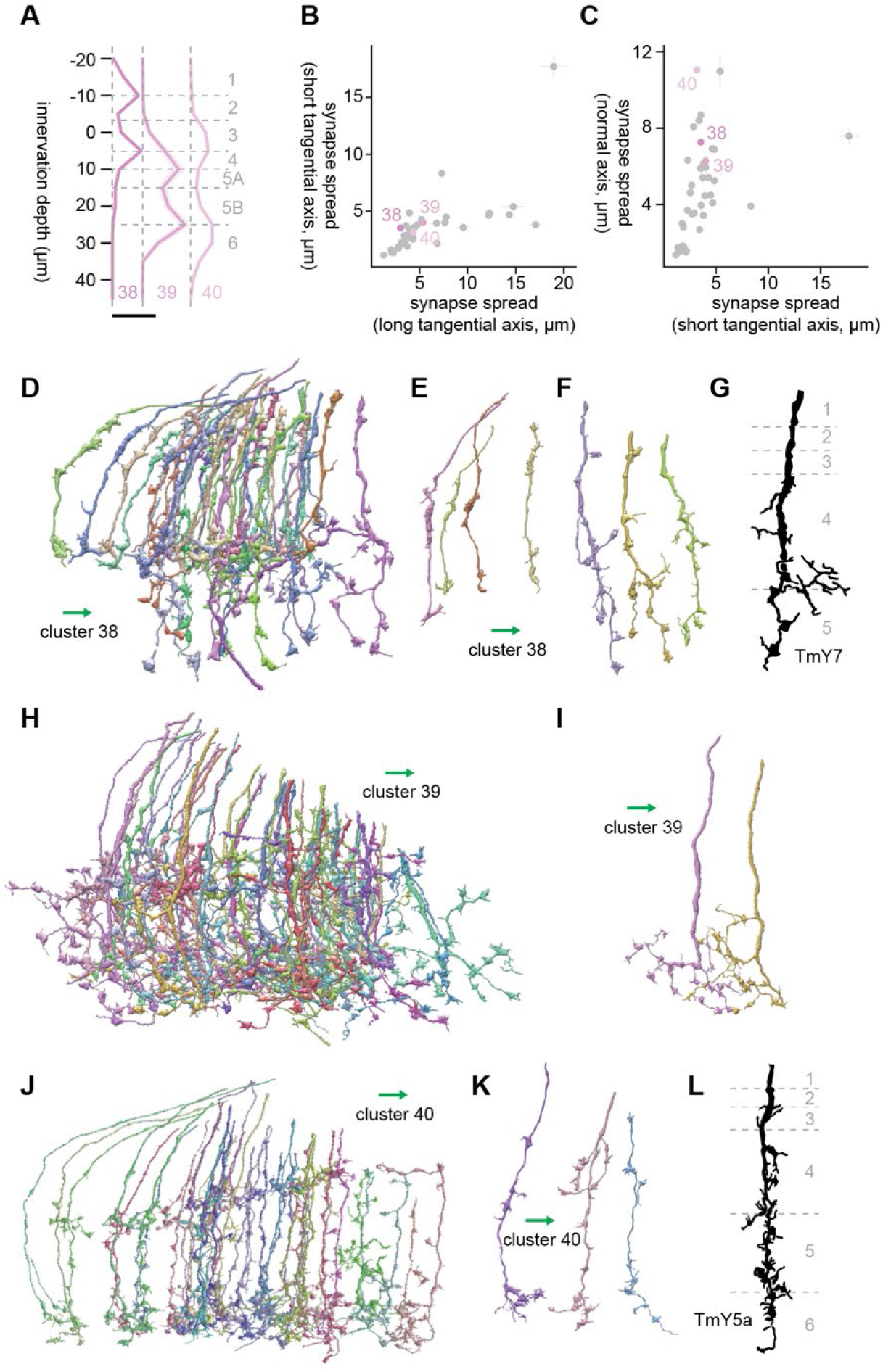
Morphology of clusters 38 through 40. (A) Mean innervation depth and (B, C) synapse spread along the three axes of clusters 38 through 40. Error bars indicate standard error of the mean. (A) Vertical dotted line indicates zero synapses. The thick horizontal bar indicates 20 synapses. (D) A population of cluster 38 cells. (E, F) Representative examples of two cell types included in cluster 38. Cells in (E) resembled cluster 6. (G) Morphology of a TmY7 axon terminal, from Fischbach & Dittrich (1989). (H) The entire population of cluster 39 cells. (I) Representative examples of cluster 39 cells. (J) A population of cluster 40 cells. (K) Representative examples of cluster 40 cells. (L) Morphology of a TmY5a axon terminal, from Fischbach & Dittrich (1989). The green arrows indicate the anterior direction.

